# Genomic analysis of natural intra-specific hybrids among Ethiopian isolates of *Leishmania donovani*

**DOI:** 10.1101/516211

**Authors:** James A. Cotton, Caroline Durrant, Susanne U. Franssen, Tesfaye Gelanew, Asrat Hailu, David Mateus, Mandy J. Sanders, Matthew Berriman, Petr Volf, Michael A. Miles, Matthew Yeo

**Affiliations:** Wellcome Sanger Institute, Wellcome Genome Campus, Hinxton, United Kingdom; Department of Parasitology, Faculty of Science, Charles University, Prague, Czech Republic; Faculty of Medicine, Addis Ababa University, Addis Ababa, Ethiopia; Faculty of Infectious and Tropical Diseases, London School of Hygiene and Tropical Medicine, London, United Kingdom

## Abstract

Parasites of the genus *Leishmania* (Kinetoplastida: Trypanosomatidae) cause widespread and devastating human diseases, ranging from self-healing but disfiguring cutaneous lesions to destructive mucocutaneous presentations or usually fatal visceral disease. Visceral leishmaniasis due to *Leishmania donovani* is endemic in Ethiopia where it has also been responsible for major epidemics. The presence of hybrid genotypes has been widely reported in surveys of natural populations, genetic variation reported in a number of *Leishmania* species, and the extant capacity for genetic exchange demonstrated in laboratory experiments. However, patterns of recombination and evolutionary history of admixture that produced these hybrid populations remain unclear, as most of the relevant literature examines only a limited number (typically fewer than 10) genetic loci. Here, we use whole-genome sequence data to investigate Ethiopian *L. donovani* isolates previously characterised as hybrids by microsatellite and multi-locus sequencing. To date there is only one previous study on a natural population of *Leishmania* hybrids, based on whole-genome sequence. The current findings demonstrate important differences. We propose hybrids originate from recombination between two different lineages of Ethiopian *L. donovani* occurring in the same region. Patterns of inheritance are more complex than previously reported with multiple, apparently independent, origins from similar parents that include backcrossing with parental types. Analysis indicates that hybrids are representative of at least three different histories. Furthermore, isolates were highly polysomic at the level of chromosomes with startling differences between parasites recovered from a recrudescent infection from a previously treated individual. The results demonstrate that recombination is a significant feature of natural populations and contributes to the growing body of evidence describing how recombination, and gene flow, shape natural populations of *Leishmania*.

**Author Summary:** Leishmaniasis is a spectrum of diseases caused by the protozoan parasite *Leishmania*. It is transmitted by sandfly insect vectors and is responsible for an enormous burden of human suffering. In this manuscript we examine *Leishmania* isolates from Ethiopia that cause the most serious form of the disease, namely visceral leishmaniasis, which is usually fatal without treatment. Historically the general view was that such parasites reproduce clonally, so that their progeny are genetically identical to the founding cells. This view has changed over time and it is increasingly clear that recombination between genetically different *Leishmania* parasites occurs. The implication is that new biological traits such as virulence, resistance to drug treatments or the ability to infect new species of sandfly could emerge. The frequency and underlying mechanism of such recombination in natural isolates is poorly understood. Here we perform a detailed whole genome analysis on a cohort of hybrid isolates from Ethiopia together with their potential parents to assess the genetic nature of hybrids in more detail. Results reveal a complex pattern of mating and inbreeding indicative of multiple mating events that has likely shaped the epidemiology of the disease agent. We also show that some hybrids have very different relative amounts of DNA (polysomy) the implications of which are discussed. Together the results contribute to a fuller understanding of the nature of genetic recombination in natural populations of *Leishmania*.

## Introduction

*Leishmania* is a diverse genus of kinetoplastid protozoan parasites from the family trypanosomatidae. These parasites are best known as the cause of human and animal leishmaniasis, which is a clinically important neglected tropical disease affecting millions of people and causing a tremendous burden of mortality and morbidity (Herricks et al. 2017; Alvar et al. 2012). Leishmaniasis comprises a spectrum of related diseases which, depending on the species, results in various presentations ranging from small, self-healing cutaneous lesions to widespread disseminated lesions, destructive mucosal and mucocutaneous pathology, and visceral disease that is usually fatal in the absence of effective chemotherapy (Herwaldt 1999). *Leishmania* have a digenetic (two host) life cycle involving a vertebrate host and 166 different species of phlebotomine sand fly that have been implicated as vectors (Akhoundi et al. 2016), although alternative invertebrate vectors may exist for some species (Seblova et al. 2015). Vertebrate hosts encompass a wide range of mammals or reptiles, and around 20 species of *Leishmania* have been reported to infect humans (Akhoundi et al. 2016).

Historically, the population structure of *Leishmania*, other trypanosomatids and indeed most protozoan parasites was considered to be largely clonal (Tibayrenc, Kjellberg, and Ayala 1990): the presumption was that admixture between members of the same clone, or between very closely related parasites was absent or rare, with minimal impact on population structure. However, at the time the clonal theory was first proposed, most population genetic data for trypanosomatids was based on inadequate sampling and use of low-resolution markers unlikely to detect admixture between genetic groups (Ramírez and Llewellyn 2014). Subsequently, extensive work using multilocus sequence typing and microsatellite markers has produced a foundation for understanding of the population genetics of some *Leishmania* species (Akhoundi et al. 2017; G. Schönian, Kuhls, and Mauricio 2011; Gabriele Schönian, Cupolillo, and Mauricio 2012; Ramírez and Llewellyn 2014). Most natural *Leishmania* isolates have surprisingly little heterozygosity, which has been widely ascribed to extensive selfing (Rougeron et al. 2009, 2010; Kuhls et al. 2007), although aneuploidy variation could also contribute (Sterkers et al. 2014). In contrast there have also been a number of reports of heterozygous natural isolates possessing a mixture of alleles associated with different populations (Schwenkenbecher et al. 2006; Kuhls et al. 2013; Gelanew et al. 2014) and even different species (Ravel et al. 2006; Hamad et al. 2011), suggesting a hybrid origin of these isolates. There is therefore a growing body of evidence for genetic exchange between natural populations of several *Leishmania* species.

Laboratory genetic crosses between at least two *Leishmania* species have been achieved in the sand fly vectors (Akopyants et al. 2009; Sadlova et al. 2011), and viable hybrids have been achieved between geographically disparate sources of *L. major* (Inbar et al. 2013). Here many hybrids possess genotypes consistent with classical meiosis; however, aneuploidy with recurrent triploidy and loss of heterozygosity (LOH) were also observed. Interspecific *L. major/L. infantum* crosses have also been performed with segregation of cutaneous and visceral traits (Romano et al. 2014). However, distinct male or female gametes of *Leishmania* have not been described, although haploid stages of *Trypanosoma brucei* have recently been discovered in tsetse flies (Peacock et al. 2014).

The fact that *Leishmania* can undergo genetic exchange is of profound epidemiological importance. Genetic exchange may facilitate adaptation to new vectors, mammalian hosts or other ecological niches. For example, *Leishmania infantum/major* hybrids infect *Phlebotomus papatasi*, a non-permissive vector for *L. infantum* that is widespread in the Indian subcontinent (Volf et al. 2007). Hybrids between *L. braziliensis* and *L. peruviana* have also been implicated as agents of destructive forms of mucocutaneous leishmaniasis (Nolder et al. 2007). Genetic exchange could also lead to the spread between populations of genes associated with resistance to drugs. Reassortment can potentially affect sensitivity and specificity of diagnostic methods and hybrid vigour (heterosis) could also affect virulence or transmission potential. Such implications are particularly worrying in the context of recombination contributing to the generation of novel visceralising traits in populations previously causing only dermal symptoms, or if adaptation to new vector species allows existing visceralising parasites to become more widespread.

Genome-wide sequence data are crucial to explore fully the extent of hybridisation and to identify the mechanisms by which hybrids are formed (Twyford and Ennos 2011). Data from only a few genetic loci may be adequate for identifying hybrids if admixture is recent or the populations have not extensively interbred. However, sparse markers are less sensitive in identifying complex, infrequent or ancient admixture, where only a small fraction of the genome may derive from any one parent. These kinds of events are known to occur in microbial eukaryote pathogens (Ropars et al. 2018; McMullan et al. 2015; Desjardins et al. 2017). Being able to describe signatures of genetic exchange in detail is important as other processes can explain hybrid patterns of genotypes; for instance, parasexual processes based on fusion of cells followed by mitotic crossing-over have previously been observed in some protozoa and are well-described in fungi (Ene and Bennett 2014). Parasexual recombination can also produce similar inheritance patterns and some evidence of this is seen in an experimental cross between different *Leishmania* species (Romano et al. 2014), where many progeny are also highly aneuploid. Whole-genome sequence data have been reported for only a single population of hybrid *Leishmania* from Turkey (Rogers et al. 2014). This population appears to have originated from a single hybridisation event between genetically disparate lineages within the *L. donovani* species complex. One of the parents appeared to be an *L. infantum*, but the precise parentage of this population remains unclear as no parental genotypes were isolated in the same region. While genomic patterns were consistent with meiosis they do not formally exclude the possibility of a parasexual process.

Here we present a detailed genomic analysis of a natural hybrid population of *L. donovani* originating from Ethiopia. East African strains of *L. donovani* are particularly diverse, consisting of two main populations: one comprising strains from northern Ethiopia and Sudan, the other strains from southern Ethiopia and Kenya (Zackay et al. 2018) and correspond to the areas populated by two different major sand fly vectors – *Phlebotomus orientalis* being the main vector in northern Ethiopia and Sudan and *Phlebotomus martini* in the South – although other vectors have also been implicated (Seblova et al. 2013). These two geographically (and genetically) isolated populations of *L. donovani* in Ethiopia also differ in clinical phenotypes (Gelanew et al. 2010). High inbreeding, seemingly incompatible with strict clonality, was observed in strains from northern Ethiopia. Microsatellite (Gelanew et al. 2010) and MLST markers (Gelanew et al. 2014) have confirmed the presence of sympatric putative parental genotypes and hybrid progeny genotypes of *L. donovani* in isolates from the northern population. Here, we apply whole-genome sequencing data to characterise more fully these Ethiopian *L. donovani* to confirm that isolates are true hybrids that originate from recombination between two different sympatric lineages. We reveal a complex pattern of inheritance implying multiple independent origins from similar parents, and backcrossing with parental types. Extensive polysomy, at the level of chromosomes, is apparent in some hybrids, the significance of which is discussed.

## Results

### Genome sequencing

From each of 11 putative hybrid isolates, 1,600-2,300 Mb of sequence data (llumina 100 bp paired-end reads) (Table 1) were generated. When mapped against the reference genome for Ethiopian *L. donovani* (Isolate LV9, WHO code: MHOM/ET/67/HU3, Rogers et al. 2011) these data produced at least 40-fold median coverage across the isolates. Generally, coverage was consistent across the genome, with more than 30 reads per sample covering at least 90% of the genome (Table 1). SNP calling identified an average of 75,775 SNPs between each individual isolate and the reference genome, and this was relatively consistent across the panel, varying between 63,042 and 89,636 (Table 1). In contrast to this consistency in the number of variable sites, the proportion inferred to be heterozygous varied considerably. Some isolates showed very low heterozygosity (for example LdM256, 0.015) while for others almost half of variant sites were inferred to be heterozygous (for example LdDM62, 0.468). Most isolates exhibiting low levels of heterozygosity (<0.1) were previously identified as putative parental genotypes (Gelanew et al. 2014), in contrast putative hybrids showed much higher levels (>0.3); with the exception of one putative parental type (LdDM481) with heterozygosity more similar to that of parental isolates (0.267). In terms of large (>100 bp) structural variants, we observed 368 deletions, 282 inversions, 169 duplications and 264 translocations. However, many of these variants do not segregate among the recent Ethiopian isolates sequenced here (see Figure S1), with most being heterozygous in all or most of these isolates (Table S1, Figure S1). A single 18 bp homozygous insertion on chromosome Ld33 was present in all of the Ethiopian isolates sequenced here, which was also present in reference strains LV9 and JPCM5 but not present in BPK282.

**Table 1:** Sequencing data and summary statistics of read mapping and variant calls.

| isolate | WHO name | Previous classification (Gelanew et al, 2014) | Sequence reads (total Mb) | Median mapped coverage | Proportion of genome with $\geq 30\times$ coverage | # of variable sites against LV9 | Heterozygosity |
| --- | --- | --- | --- | --- | --- | --- | --- |
| LdDM19 | MHOM/ET/2007/DM19 | Possible hybrid* | 21383530 (2138.3) | 53 | 0.97 | 80930 | 0.301 |
| LdDM20 | MHOM/ET/2007/DM20 | parental type A | 19098154 (1909.8) | 49 | 0.96 | 74749 | 0.024 |
| LdDM62 | MHOM/ET/2007/DM62 | hybrid | 19577434 (1957.7) | 45 | 0.94 | 89636 | 0.468 |
| LdDM256 | MHOM/ET/2008/DM256 | parental type B | 19854188 (1985.4) | 51 | 0.96 | 63042 | 0.015 |
| LdDM257 | MHOM/ET/2008/DM257 | parental type B | 23066268 (2306.6) | 59 | 0.97 | 63088 | 0.016 |
| LdDM259 | MHOM/ET/2008/DM259 | parental type B | 16023002 (1602.3) | 42 | 0.91 | 63209 | 0.028 |
| LdDM295 | MHOM/ET/2008/DM295 | hybrid | 20010052 (2001.0) | 51 | 0.96 | 87546 | 0.388 |
| LdDM297 | MHOM/ET/2008/DM297 | parental type A | 20651456 (2065.1) | 52 | 0.96 | 76530 | 0.072 |
| LdDM299 | MHOM/ET/2008/DM299 | hybrid | 22442162 (2244.2) | 53 | 0.97 | 88605 | 0.462 |
| LdDM481 | MHOM/ET/2009/DM481 | parental type B | 20625354 (2062.5) | 53 | 0.97 | 82728 | 0.267 |
| LdDM559 | MHOM/ET/2009/DM559 | parental type B | 18753496 (1875.3) | 51 | 0.96 | 63469 | 0.030 |

### Genome-wide variation identifies distinct subgroups with different levels of heterozygosity

A phylogenetic tree and principal components analysis of the SNP data suggesting that these 11 isolates are divided into multiple groups, with two parental groups and at least two putative hybrid groups (Figure 1). As expected the two parental groups are composed of isolates with low heterozygosity. The first parental group, comprises LdDM259, LdDM559, LdDM257, LdDM256 and the second, comprises LdDM20, LdDM297. LdDM481 is an outlier in that it is a highly heterozygous sample that forms a distinct lineage, intermediate between the two parental groups but clearly distant from the other four putative hybrid isolates (LdDM19, LdDM62, LdDM295, LdDM299). Two isolates (LdDM62 and LdDM299) isolated from the same patient (a post treatment recrudescence in an HIV patient) appear very similar on the tree and PCA. The first two principal components (PCs; Figure 1b) explain 86.32% of the variance (60.32% and 20% respectively) in the data broadly reflecting previous interpretation of these isolates as hybrids and parental genotypes, the putative hybrid isolates being intermediate between the sets of parentals. An interesting exception is LdDM481, which appears distinct from all other samples regarding the first PC, with all subsequent PCs showing similar patterns, up to PC5, where LdDM19 appears as distinct from the other isolates: however, this axis encompasses only 1.9% of the total variation in these data. In the phylogeny, inclusion of three additional reference genomes (LV9, an *L. donovani* isolate from an Ethiopian VL patient; JPCM5, a Spanish canine *L. infantum*, and BPK282 from a Nepalese VL case) revealed the diversity present in this Ethiopian cohort and their distant relationship to both *L. donovani* in the Indian subcontinent and *L. infantum* (Figure 1a). The reference isolate LV9, originally isolated in 1967, appears to be closely related to one of the parental populations.

**Figure 1.**
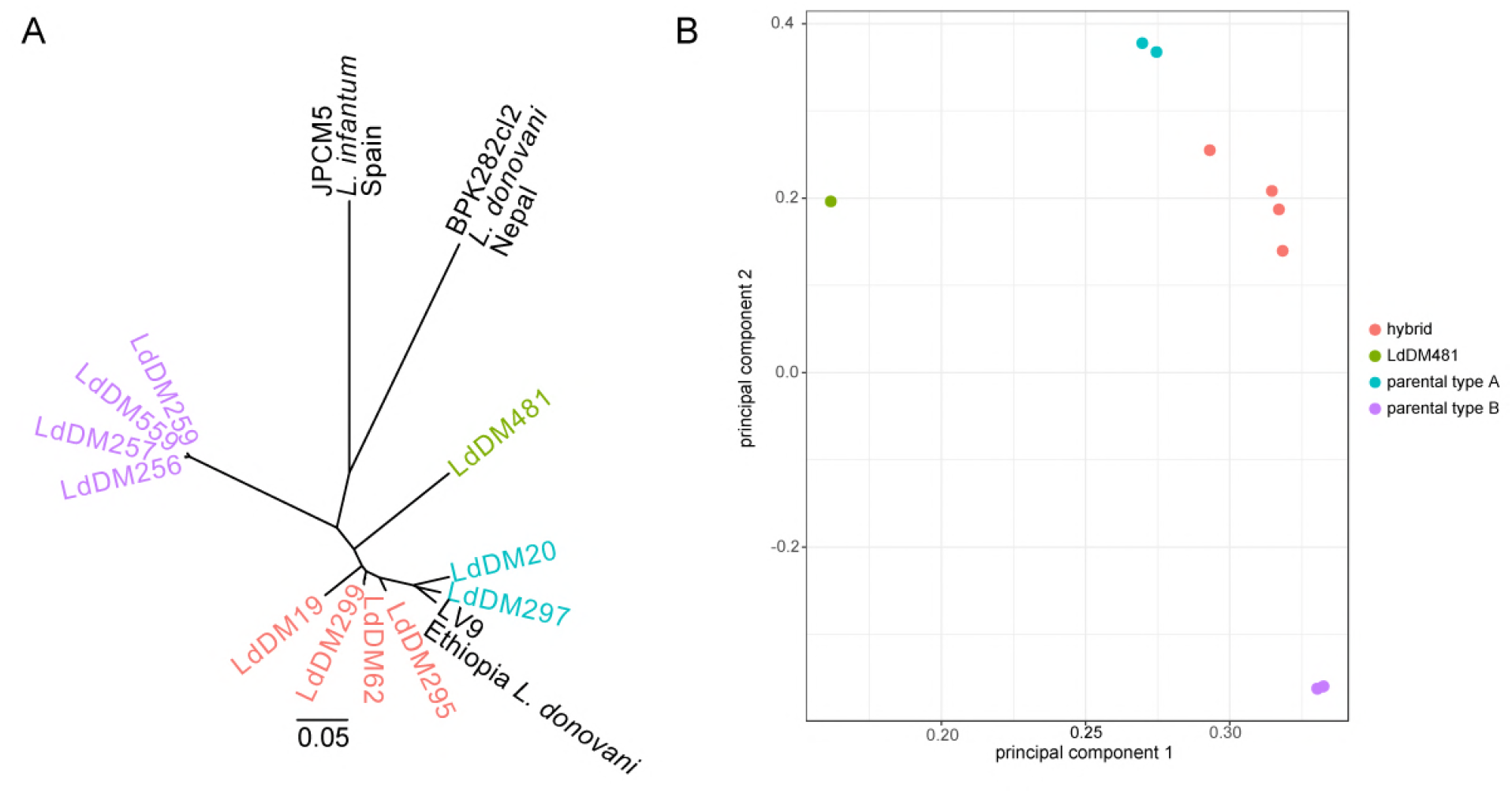
Neighbour-joining phylogeny (A) and principal components analysis (PCA; B) based on genome-wide SNP variation data among Ethiopian *L. donovani* isolates, including additional isolates from the *L. donovani* species complex. Scale bar on (A) represents genetic distances in terms of expected substitutions per nucleotide site.

Chromosome copy number for each isolate was inferred from read depth and allele frequencies at heterozygous sites (see methods, Figure 2). Broadly, chromosomes across the majority of isolates were inferred to be diploid. The exception is chromosome 31, which, as usual in *Leishmania*, was inferred to be highly polysomic with at least four copies present in all isolates. Chromosomes 5, 6, 8, 20 and 35 were also observed at higher dosage, being at least trisomic in 6 of the 11 isolates. Two samples stood out as being more highly polysomic than others: LdDM19 was inferred to be tetrasomic at three chromosomes (13, 31 and 3), while LdDM299 was strikingly polysomic. For this isolate, allele frequency data suggested a minimum of tetrasomy across chromosomes, with half the chromosomes inferred at even higher dosage (6 pentasomic, 9 hexasomic, 1 heptasomic, with chromosomes 31 and 33 octasomic).

**Figure 2.**
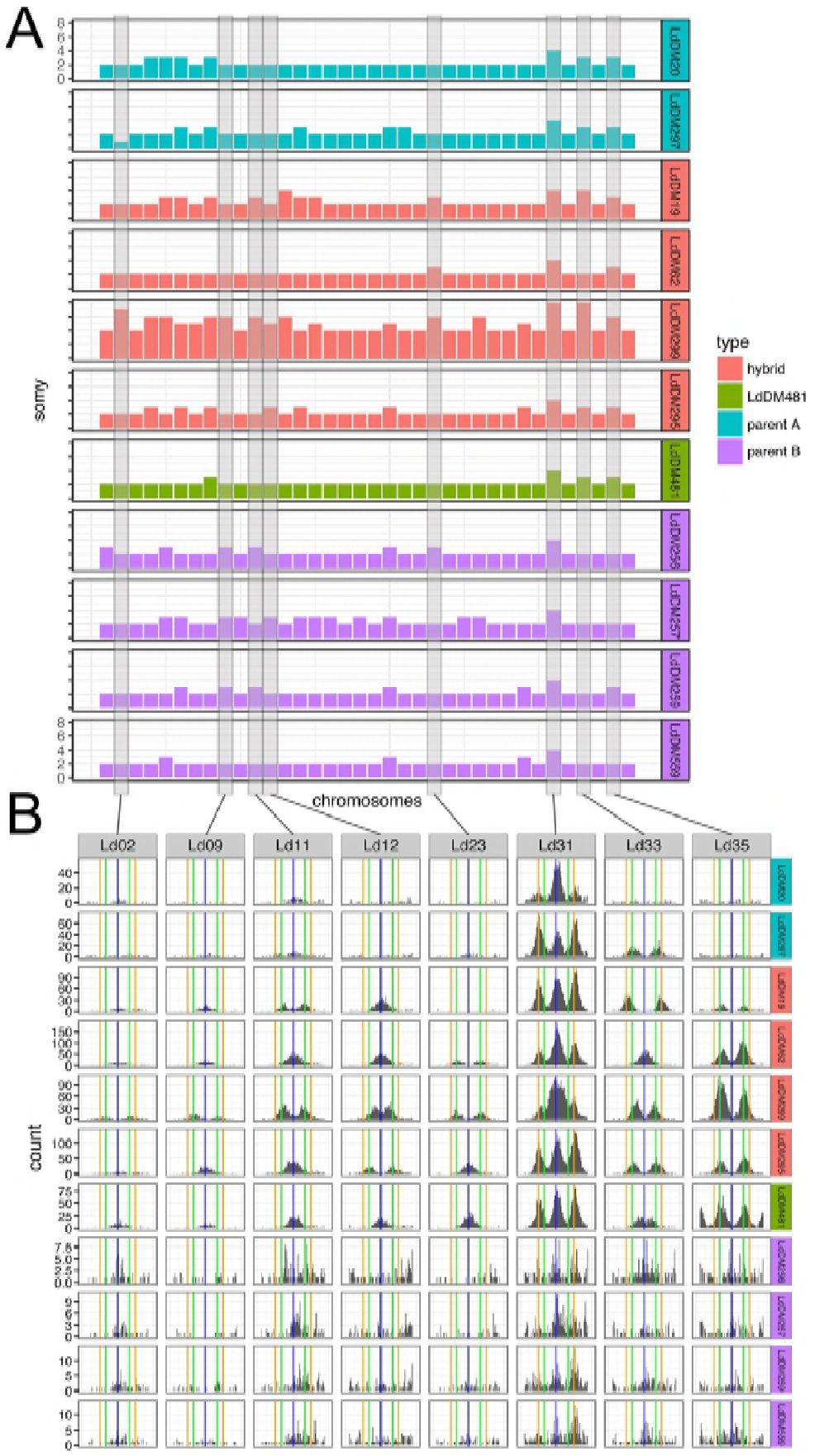
Variable somy across Ethiopian isolates inferred from coverage and allele frequencies. (A) Shows the inferred chromosome copy number (somy) for each chromosome across Ethiopian isolates under study. Y-axis scales are the same across all panel A rows. As detailed in methods, relative somy is inferred from the coverage depth of reads mapped to each chromosome, while the baseline somy is determined from the allele frequency distribution. (B) Shows example distributions of non-reference allele frequencies for each isolate, highlighting differences in somy. Vertical lines are at allele frequencies of 0.5 (blue), 0.33 and 0.67 (green), 0.25 and 0.75 (orange); expected for disomic, trisomic and tetrasomic chromosomes respectively.

The high somy of LdDM299 is of particular interest given that LdDM62, the pre-treatment sample from the same HIV infected patient, has somy similar to the other hybrid isolates. These two isolates are otherwise genetically very similar, differing at only 4,484 sites across the genome (LdDM62 differs from the other hybrid isolates at 38,023 and 25,765 sites). The difference in allele frequency distribution between LdDM62 and LdDM299 is clear: for example, chromosomes 11, 12 and 33 all show a clear peak in allele frequencies close to 0.5 in LdDM62 suggesting disomy (or at least an even chromosome dosage), but peaks at 0.33 and 0.67 in LdDM299, suggest a higher dosage of at least 3 copies (Figure 2b). The very high somy of other chromosomes is then inferred from the ratio of coverage (Figure 2a). We attempted to confirm the high ploidy of LdDM299 using flow cytometry to measure DNA content. Cells from the same population of cells that was sequenced were not available, but a cloned population separated from the sequenced cells by more than 8 *in vitro* passages was analysed. DNA content of these cells was suggestive of diploidy (S1 Fig), leaving some uncertainty about the precise somy of the LdDM299 isolate. During SNP-calling, the copy number of individual chromosomes is specified. We thus confirmed that our main results are insensitive to the assumed somy of the isolates by repeating most analyses with genotypes called as though all isolates are diploid: in all cases the conclusions from our analyses are qualitatively the same with diploid genotypes, or genotypes called using the inferred somies.

### Patterns of inheritance from putative parental populations

Variants at many sites were shared by putatively hybrid isolates and either one or other of the parental group; with relatively few variants present only in the hybrids and not found elsewhere (Figure 3). LdDM481 was an exception, possessing a moderately high number of private variants and also sharing some different variants with the parental groups compared to other hybrids, particularly with parent B where other hybrid isolates shared substantially more variation.

**Figure 3.**
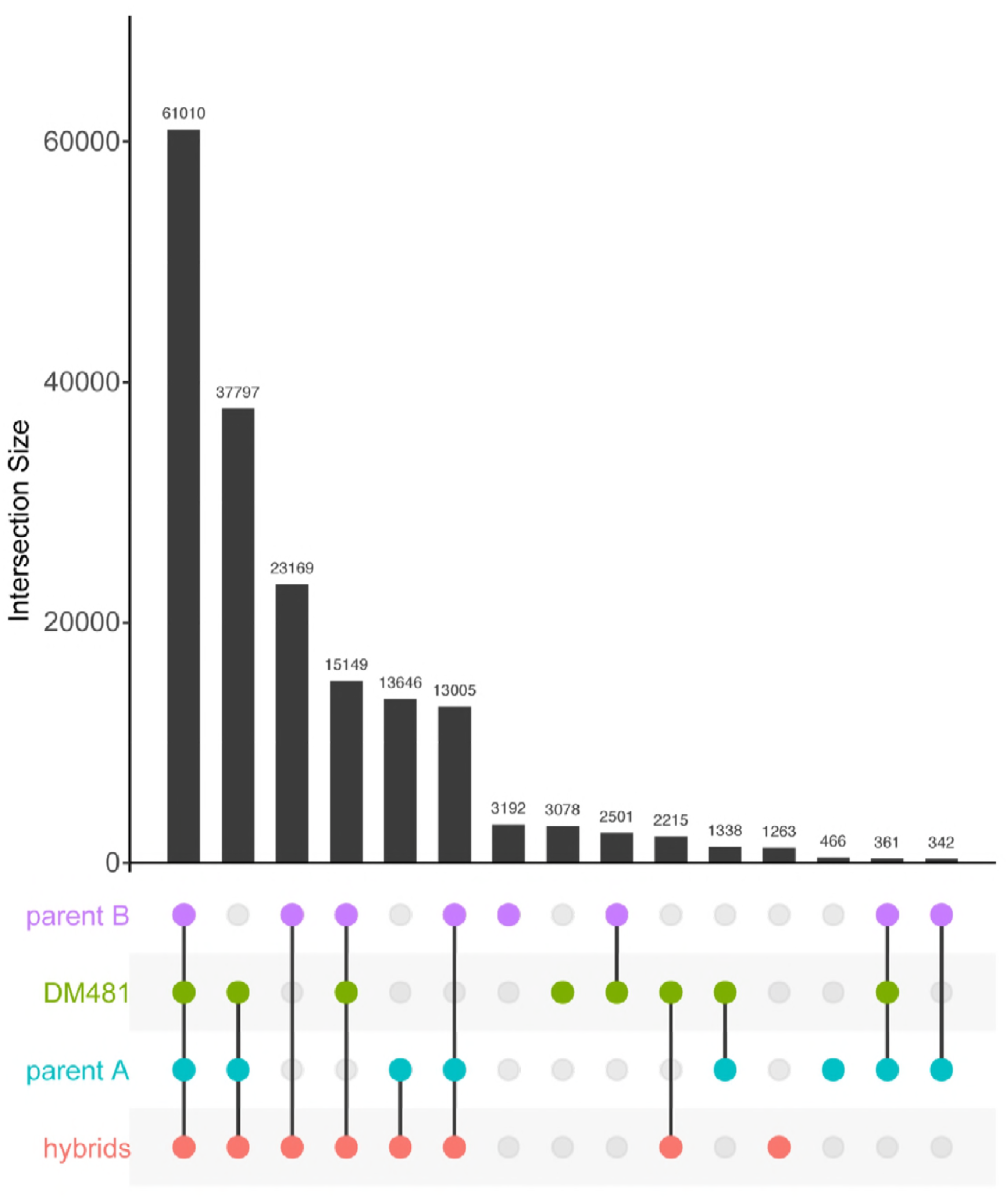
Pattern of allele sharing between groups of Ethiopian *L. donovani* isolates. Each row corresponds to one of the categories of isolates, with columns corresponding to non-reference alleles present in two categories or more (intersections), and the bar graph depicts the number of alleles present in at least one of the isolates in a group for each intersection.

To understand further the origins of the hybrid parasites, we identified SNP variants that are fixed differences between the two groups of parental parasites (excluding LdDM481) and used these as markers to identify the likely origin of variants identified in the hybrids, effectively ‘painting’ the hybrid isolate chromosomes by their likely ancestry under the hypothesis that these isolates originated as hybrids between the parental groups or close relatives. We identified a total of 49,835 such ‘parent-distinguishing SNP’ sites at which the two parental populations were completely fixed for different alleles. These sites are distributed across all chromosomes. For the vast majority of sites, across all putative hybrid isolates, genotypes consisted of a combination these parental alleles (Table 2), supporting the notion that these isolates originated as hybrids between two parental types. The different categories of sites are strikingly unevenly distributed across the genomes suggestive of genome wide hybridisation (Figure 4). For some chromosomes these SNP markers appear to be uniformly inherited from a single parent or show uniform heterozygosity with one allele from each parental type. Most chromosomes, however, show multiple blocks of sequential SNPs of different origins, revealing a patchwork of blocks of different ancestries. Isolate LdDM481 is an exception: the “blocky” structure visible in other isolates is not apparent, there are less than half the number of heterozygous sites of shared origin in comparison to any other hybrid isolates, and five times the number of genotypes containing alleles other than those fixed in the parental groups, although this still represented only 54 sites.

**Figure 4.**
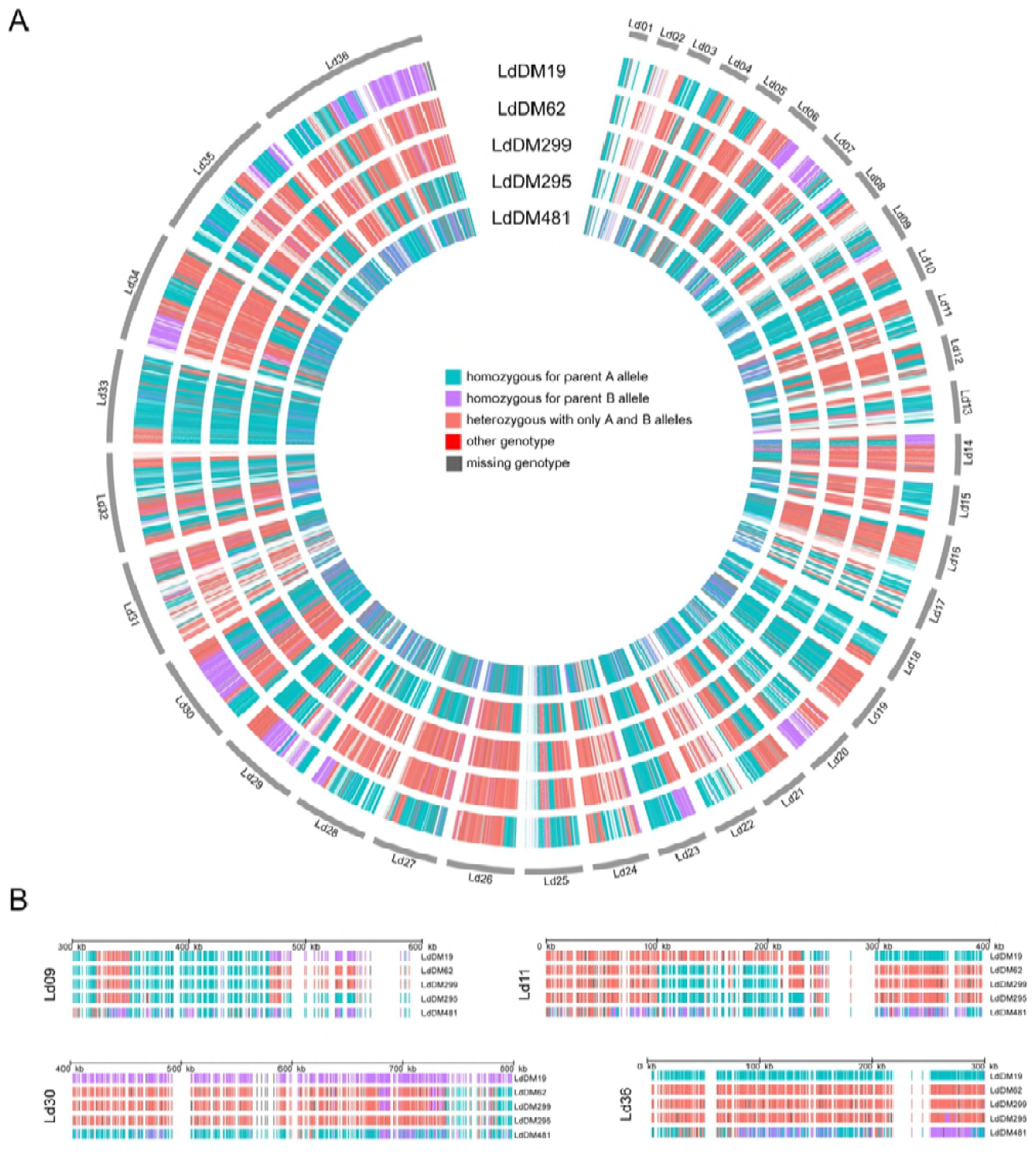
Distribution of SNPs distinguishing between potential parents. (A) Coloured bars in concentric rings represent every SNP that are fixed homozygous differences between the two sets of putative parents, colored to represent the diploid genotype call as homozygous for either parental type or heterozygous with one of each allele. A small number of sites had other genotypes or no reliable genotype call. There are very few SNPs in the red or grey categories. (B) Shows a magnified view of the same data for four regions, chosen to highlight variation between different isolates.

**Table 2.** Distributions of parental-distinguishing SNPs in putative hybrid isolates. Values are numbers of sites in each isolate from each category. For polysomic chromosomes, sites with at least one allele from each parent are grouped as ‘heterozygous AB’ irrespective of dosage of A and B alleles.

| isolate | Homozygous parent A | Homozygous parent B | Heterozygous AB | Genotype not unambiguously inferred from reads | Other genotype |
| --- | --- | --- | --- | --- | --- |
| LdDM19 | 21,171 | 10,419 | 16,633 | 1,602 | 10 |
| LdDM62 | 15,600 | 2,103 | 29,779 | 2,343 | 10 |
| LdDM295 | 25,482 | 1,429 | 21,032 | 1,880 | 12 |
| LdDM299 | 15,868 | 2,036 | 18,571 | 13,352 | 8 |
| LdDM481 | 31,104 | 10,271 | 7,400 | 1,006 | 54 |

To get a more quantitative understanding of the distribution of these parent distinguishing SNP sites in blocks inherited from each parent we developed a hidden Markov model (HMM) to assign putative ancestry for sites without parent-distinguishing SNPs. The HMM assigns every 100 bp window of the genome to three states: either homozygous ‘parent A’, homozygous ‘parent B’ or heterozygous (Figure 5a). The advantage of the HMM is that it can statistically assign windows even in the absence of any parent-distinguishing variants (or where those variants are ambiguous), under the assumption that transitions between the states occur in a regular way across the genome if there is no direct evidence of a change. The HMM assigns significantly different proportions of the genome to each category for different isolates (Figure 5a), and the uneven distribution of sites in each category across the genome is reflected in the fact that these proportions are quite different from the proportion of parent-distinguishing SNP sites assigned to each category. This model also estimates the length of ‘runs’ that form blocks of genome with a single inheritance pattern. The distribution of these block lengths provides information on the relative age of hybrids, as we would expect recombination to have broken down blocks from older introgression events more than recent events. Longer blocks suggest that fewer hybridisation events have occurred between admixed clones but backcrossing to parental clones or related parasites would also contribute to longer blocks. The block length distribution varies between isolates (Figure 5b): it suggests that LdDM62 and LdDM299 represent a more recent hybridisation with a ‘parent B’ type than any ‘parent A’ type, and that LdDM295 may originated from a more recent hybrid (or with fewer hybridisations) between the two parental types than LdDM19. For at least two isolates (LdDM62 and LdDM299) the inheritance is asymmetrical, in that they are inferred to have inherited different proportions of their genome from each of the parental types, suggesting that these isolates did not originate by crossing within a ‘founder’ hybrid population, but involved some degree of backcrossing with parent B types.

**Figure 5.**
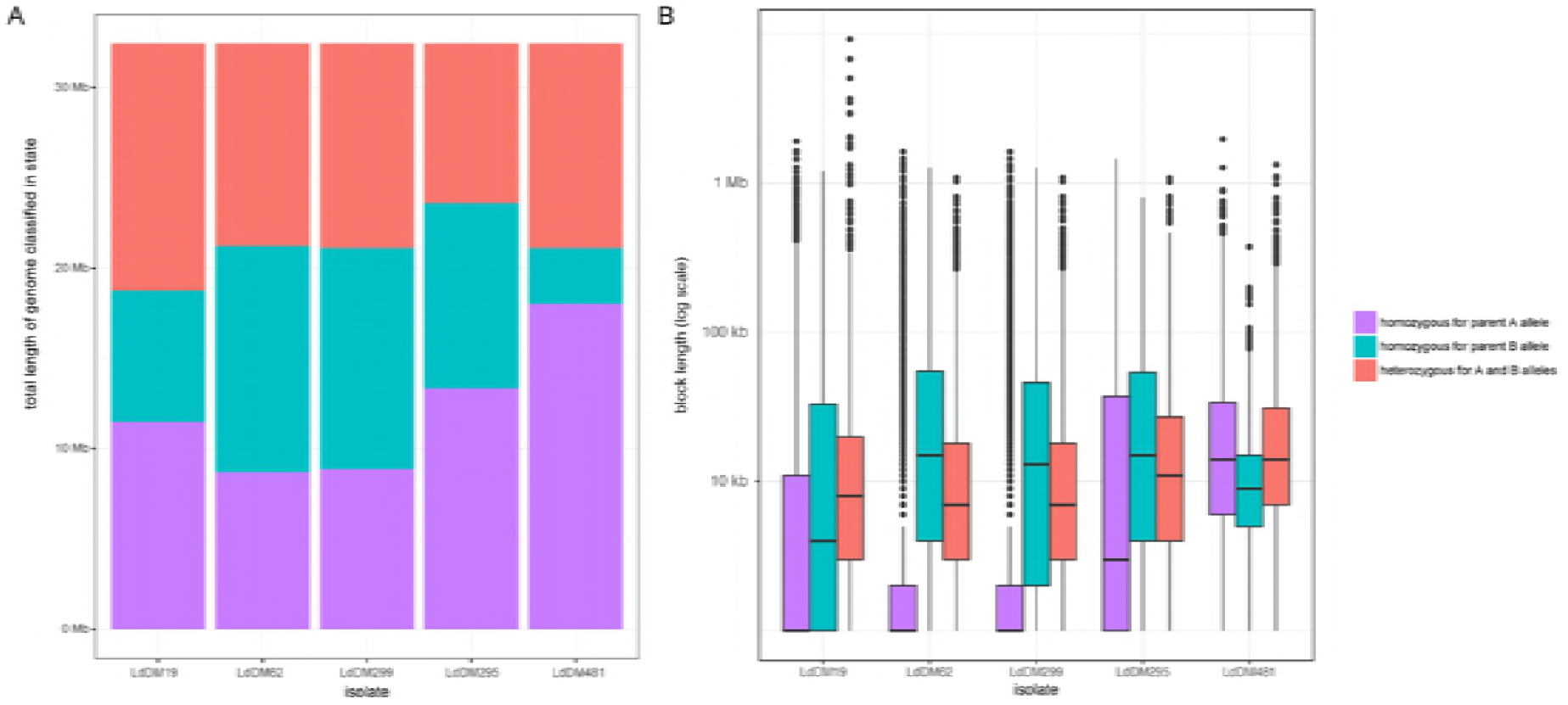
Distribution of genomic regions of putative hybrid isolates between parental origins based on Hidden Markov Model. (A) Shows the total number of basepairs assigned as homozygous parent A, homozygous parent B and heterozygous based on the maximum posterior probability assignment of hidden states of the Hidden Markov Model. (B) Box-and-whisker plot showing the distribution of lengths of contiguous blocks assigned to each of these three parentage states across the 5 putative hybrid strains in the most probable path identified in the Viterbi decoding. Boxes show median length and interquartile range on a log axis, whiskers are

### Reconstructing haplotypes confirms that variants of each parental type are linked

We used reads and read pairs to phase locally heterozygous sites that were physically close to each other in the genome into sets of variants known to be present on a single haplotype. We subsequently identified regions at which all heterozygous positions were phased in all 15 isolates, so that we have unambiguous information about the haplotypes present in these regions. We inferred haplotype phylogenies for nine such regions that were at least 3 kb long and had an average of at least 4 heterozygous sites per isolate; this included 9 of the 16 ‘fully phased’ regions of 3 kb or longer (figure 6). All 9 blocks were on different chromosomes.

**Figure 6.**
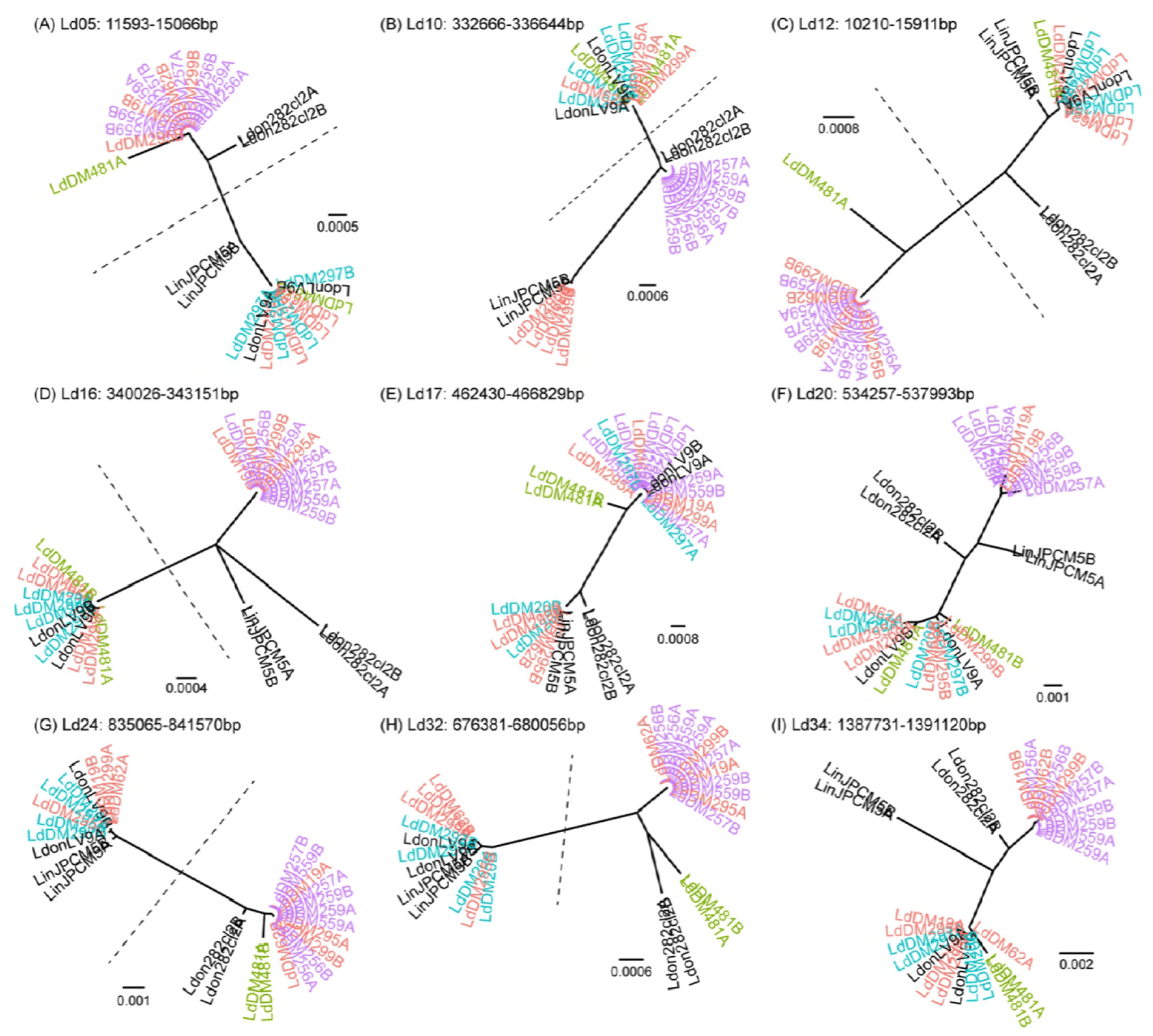
Maximum-likelihood phylogenies for inferred haplotypes at 9 genome regions at which all isolates could be phased into blocks of length greater than 3kb and with an average of at least 4 heterozygous sites per isolate. A and B labels on the leaves are arbitrary names for the two different haplotypes at each locus, for each isolate. Dotted lines separate the two hybrid haplotypes for those blocks at which all the hybrid isolates (except Ld481) have one haplotype from each putative parental population.

In most blocks (7 out of 9; Figure 6a-e, g, h) the two haplotypes for each of the four hybrid isolates (LdDM19, LdDM62, LdDM295 and LdDM299) cluster in different parts of the phylogeny. For 6 of these (Figure 6a-d, g, h), phylogenies show the expected pattern if the hybrid isolates originated from a simple, single hybridisation between the two parental types: a long branch of the haplotype tree separates all parent A haplotypes together with one haplotype of each of the 4 hybrids from parent B haplotypes with the second haplotype of each hybrid. In one block (Figure 6e) on chromosome 17 the two haplotypes for each hybrid isolate divide into two clusters as expected, but the two ‘parent A’ isolates (LdDM20 and LdDM297) appear in different clusters. In the chromosome 34 block, both haplotypes for one hybrid isolate (LdDM295) clustered with the same parental group – the parent A isolates (figure 6i). All of these haplotypes are consistent with a hybrid origin for the isolates but with a more complex history than a simple, single hybridisation between the two parental populations. This could involve further recombination either by ‘intercrossing’ within the hybrid population or backcrossing to the parental types.

Only the final block (Figure 6f) does not support a simple hybrid origin for these isolates, as the two haplotypes for each isolate cluster together. Further rounds of crossing, with a different history for LdDM19 and the other hybrid isolate, would explain the pattern at this locus. Examining the alignments for these blocks did not reveal any sign of recombination within reconstructed haplotypes, but did reveal some haplotypes in the putative hybrids that differ from either of the putative parental types - for example at the haplotype block on Ld10 all of the hybrids share one haplotype that is very similar to those in *L. infantum* JPCM5 but missing from any of the other *L. donovani* isolates (Figure 6b; figure 7a). Either the parent B isolates are a poor proxy for the true parental types at this locus, or the history of the hybrid isolates includes crossing with more than two parental populations.

**Figure 7.**
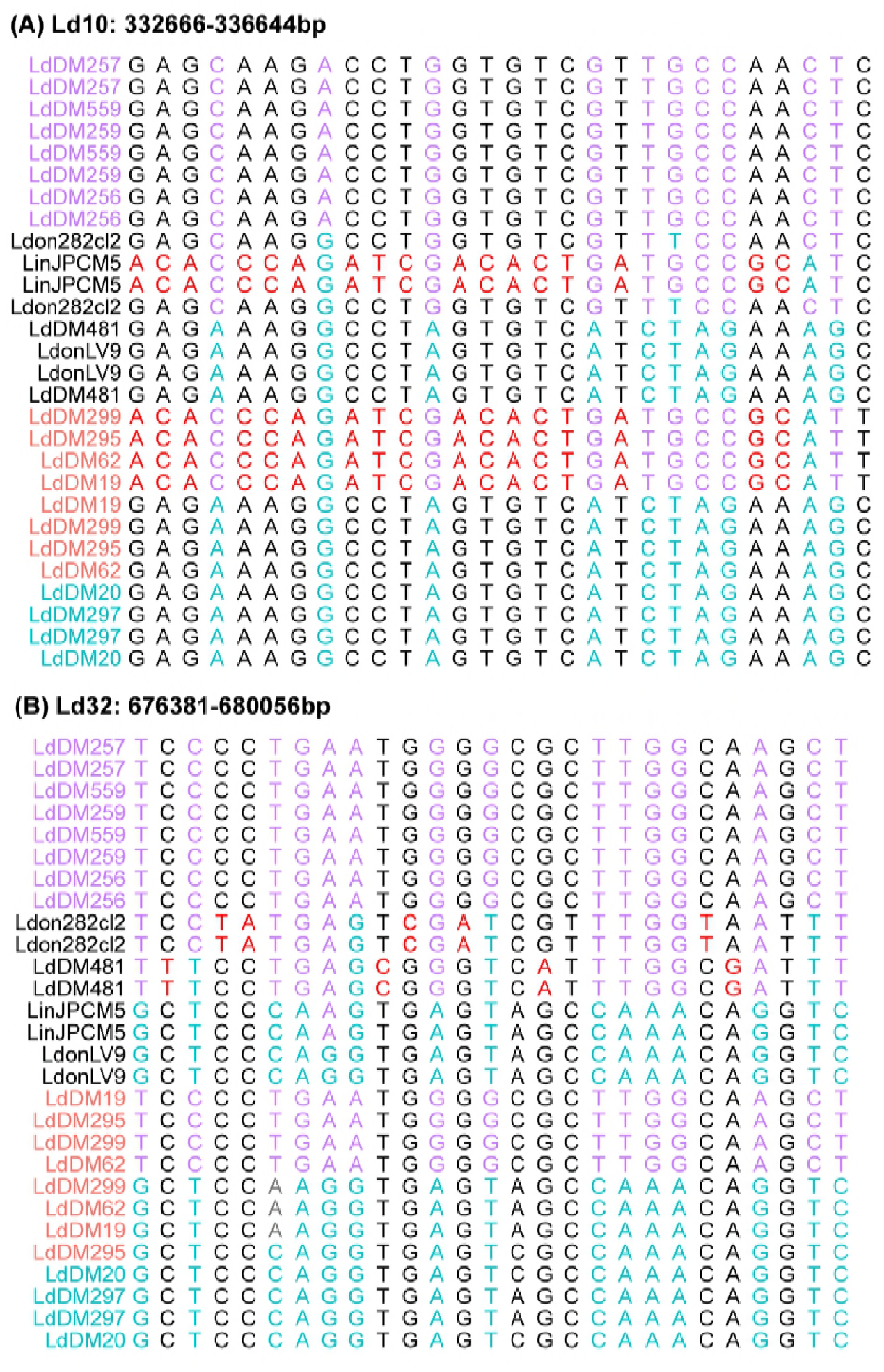
Alignments of inferred haplotypes in fully-phased regions, showing unusual haplotypes in (a) the hybrid isolates and (b) LdDM481. Panels show all variable sites within two of the ‘fully phased’ genome regions shown on figure 5. In all panels, red sites identify alleles specific to two unusual haplotypes discussed in the text. Cyan and magenta identify sites at which parent A and parent B isolates share fixed different alleles (parent distinguishing sites).

The pattern for LdDM481 haplotypes was more complex: in 7 out of 9 trees the two haplotypes for this isolate appear in the same cluster: twice with parent B haplotypes (figure 6g,h) although never very closely related to these; four times with parent A haplotypes (figure 6b,d,f,i). In one other case (Figure 6e) the parent A isolates are themselves non-monophyletic. At many of these loci, LdDM481 has haplotypes not present in other isolates. At two other loci (on Ld05 and Ld12; Figure 6a, c) LdDM481 is heterozygous for one haplotype not observed elsewhere and for one haplotype shared with parent A isolates and the hybrids. At the phased locus on Ld32 (Figure 6h), the LdDM481 haplotype is apparently distantly related to BPK282; although closer inspection of this locus shows they are united by their lack of alleles present in other isolates rather than shared characters (Figure 7b).

For the other “outgroup” isolates of *L. infantum* and *L. donovani*, the two haplotypes from each isolate consistently cluster together. Wherever parent A haplotypes are monophyletic, the Ethiopian LV9 isolate haplotypes group with them. Nepalese *L. donovani* BPK282 tends to group with the parent B isolates but is often clearly distinguishable from them; the position of *L. infantum* JPCM5 haplotypes on these trees is more variable.

### Kinetoplast (kDNA) phylogeny

The kDNA maxicircle is homologous to the mitochondrial genome of other eukaryotic groups (Jensen and Englund 2012), and is thought to be uniparentally inherited in *Leishmania* (Akopyants et al. 2009) and trypanosomes (Turner et al. 1995). The maxicircle phylogeny (Figure 8) shows a close relationship between the hybrids, parental type A, LdDM481, and the historical reference LV9 isolate, and are phylogenetically distinct from parental B isolates. This contrasts with the nuclear phylogeny, which shows the hybrid samples as somewhat more closely related to parent A isolates but clearly intermediate between both parental groups: here, the parental A isolates do not even form a monophyletic group, with LV9 and hybrid LdDM297 clustering with one parental type. Surprisingly, the mitochondrial phylogeny suggests some divergence between LdDM299 and LdDM62, despite them originating pre and post treatment from the same patient. However, supporting bootstrap values were low for all relationships in this part of the tree. The outlier isolate LdDM481 also appears less distant from the cluster comprising parental group A and hybrids than indicated from nuclear SNP variation, even though mutation rates are likely to be an order of magnitude greater for mitochondrial data than for the nuclear genome. The uncertainty in the precise relationships between isolates notwithstanding, the close relation between parental group A and the hybrid isolates suggests the hybrids uniparentally inherited parent type A mitochondrial genomes, and gives some support to the idea that these all originated from a single initial cross, although these data cannot exclude that there is some bias in the inheritance of kDNA between strains, or that this shared kDNA type reflect subsequent backcrossing rather than the original hybridisation.

**Figure 8.**
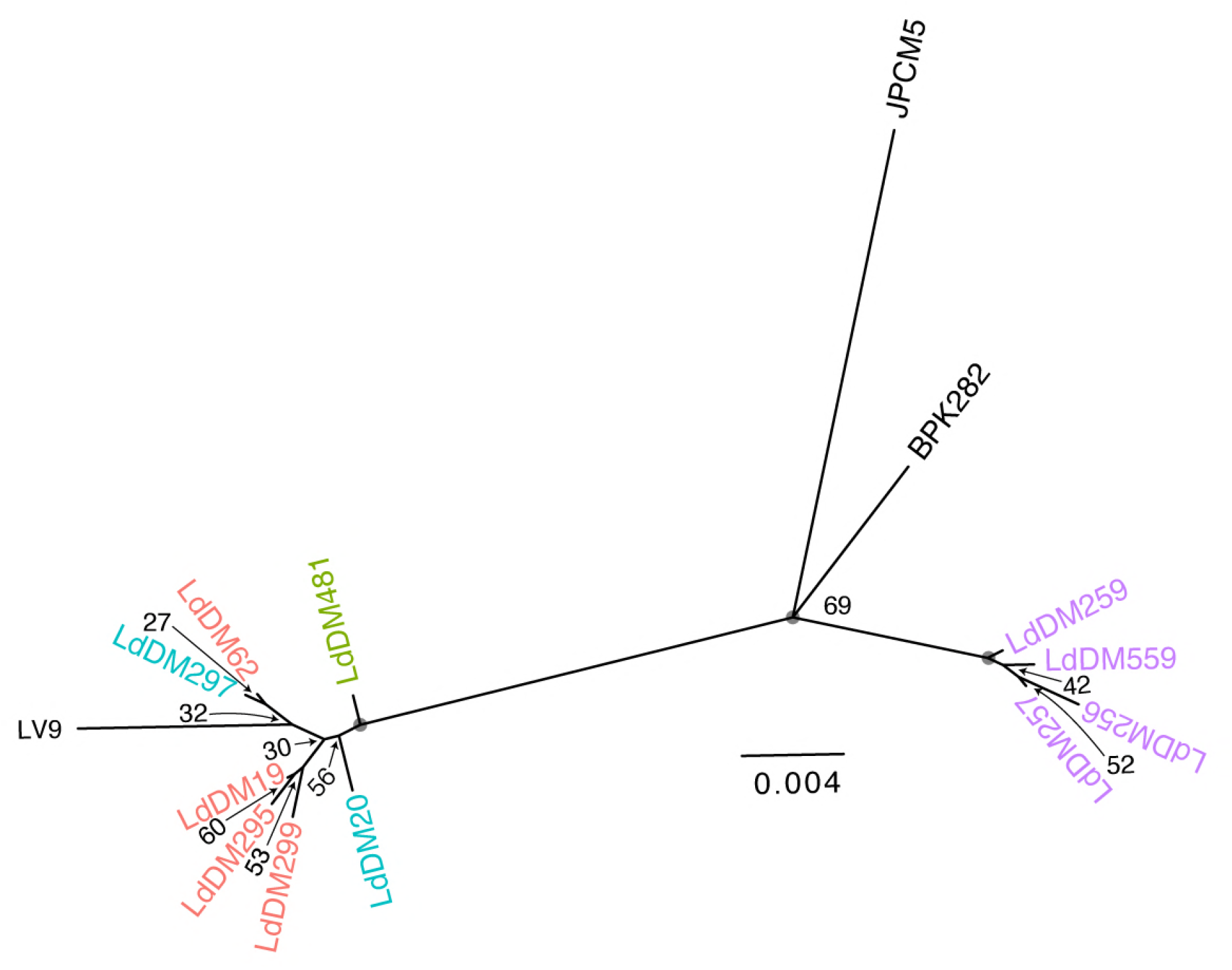
Maximum-likelihood phylogeny of reconstructed mitochondrial (kDNA maxicircle) haplotypes. Values on nodes are bootstrap support values for the partition induced by deleting the edge below each node, grey circles denote 100% support.

## Discussion

Hybridisation in *Leishmania* has now been demonstrated experimentally and is also observed in natural isolates across different species. These include multiple *L. braziliensis-L. peruviana* hybrids in Peru (Nolder et al. 2007), *L. infantum-L. major* hybrids (Volf et al. 2007) and a widespread lineage of *L. tropica* that appears to be disseminated from a recent hybridization event (Schwenkenbecher et al. 2006). However, the only previous whole genome analysis of hybrid *Leishmania* isolates identified a population from Turkey that appeared to be hybrids between a MON-1 genotype type of *L. infantum* and another member of the *L. donovani* species complex (Rogers et al. 2014). In that case, isolates appear to be generated solely from continued crossing within an initial hybrid population without back-crossing to either parental type. The genetic heritage of Ethiopian hybrids we describe here must be more complex. More specifically we propose that these isolates must have originated from more than one crossing event between similar parents. Additionally, crosses must have occurred between either different hybrids types or between hybrids and parentals subsequent to the ‘founding’ outcrossing event.

In more detail, our results are indicative of complex *L. donovani* populations in northern Ethiopia, with the relatively small number of isolates sequenced here representing at least three different histories. We find two distinct parental groups of isolates with low heterozygosities. The first parental group, comprising LdDM259, LdDM559, LdDM257, LdDM256 and the second, comprising LdDM20, LdDM297. We confirm that four of the heterozygous isolates (LdDM19, LdDM62, LdDM295 and LDM299) possess hybrid genotypes that are phylogenetically intermediate between these two parental groups. Most blocks of variants in these genomes can be interpreted as being inherited from one, or both of the parental populations. The large-scale structure of these genomes as blocks of variants of particular ancestry suggests that these are relatively recent events, at least in terms of the number of crossing-over events that have occurred since. There are however substantial differences between the hybrids in that the number of variants shared by individual hybrid isolates and the parentals, and also their distribution across the genomes differs substantially.

Results are indicative of at least three different histories of crossing between parental types, and probably within the hybrid population itself, with LdDM299 and LdDM62 showing very similar patterns, while both LdDM19 and LdDM295 are distinct. Using a hidden Markov model, we estimated the lengths of blocks of sequence inferred to originate from each parental population. The distribution of sizes of contiguous sequence fragments derived from each parent was not consistent between the hybrids, probably reflecting differences in the timing of hybridization events; we expect that continued interbreeding and the accompanying crossing-over gradually broke down these blocks of ancestry. Hence, blocks inherited from parents in older crossing events have smaller stretches of continuity. Our HMM results thus gives qualitative insight into the relative age of the different events that gave rise to these hybrid isolates. Using this approach we infer that LdDM62 and LdDM299 have crossed onto a parent B like ancestor more recently than to parent A, while the ancestry of LdDM19 and LdDM295 from both parents is approximately equally old. This approach assessing length of inherited haplotypes has been previously used to gain quantitative results into the history of human populations (Pugach et al. 2011), but parameterising this kind of approach in *Leishmania* seems challenging, given the facultative nature of sexuality in this genus, and our current lack of understanding of recombination in *Leishmania*. In particular, we do not quantitative information on the mutation rate, recombination rate or even likely generation time of *Leishmania* in vivo. In all cases a greater proportion of the genomic variation was shared with the ‘parent A’ population, suggesting a more recent common ancestry with this population or backcrossing with this population during evolutionary history. The close relatedness between the LV9 isolate, isolated in the Humera district of Ethiopia in 1967, and one of the parental groups indicates that parental like genotypes have been present in this region for extended periods of time. While the age of the hybridisation events we have described are unclear, the presence of genotypically stable “parental donors” over time may have facilitated the emergence of multiple hybrid populations. In summary, while we cannot reconstruct the precise history of this population, our data confirm that the population of *L. donovani* in Ethiopia has undergone multiple rounds of hybridisation, including more complex patterns of crossing than simple F1 hybridisation between parents or subsequent crossing within a hybrid population.

The highly heterozygous isolate LdDM481 emerged as a consistent outlier that forms a distinct lineage, intermediate between the two parental groups but also distinct from the other four putative hybrid isolates. The presence of haplotypes at a number of loci in this isolate which are not present in either parental population makes it seem unlikely that this is a very recent hybrid between the two parental groups in the current cohort. One explanation is that DM481 is simply a representative of a distinct, divergent population, which appears plausible considering the genetic diversity within *L. donovani* that has previously been identified within this region (Gelanew et al. 2010, 2014; Zackay et al. 2018). A more unlikely scenario is that DM481 has some recent common ancestry with the parent A population but less so with population B.

The multiplicity of hybrid and parental genotypes within this small sample of isolates from northern Ethiopia suggests that genetic exchange is commonplace among *L. donovani* populations transmitted by *P. orientalis*, and that resulting hybrid progeny may be widely disseminated. This complex evolution also implies that co-infection of *P. orientalis* with different *L. donovani* isolates occurs frequently, at least in this region. It is currently unknown if hybrids are most likely to emerge when a sand fly is co-infected with different *Leishmania* strains ingested in the same blood meal or subsequent feeds. For *T. brucei* there is evidence that the production of hybrid genotypes is most successful when both parental types are taken up in a single meal by the tsetse vector (Peacock, Bailey, and Gibson 2016).

The pattern of polysomy we observe across the cohort did not reflect the phylogenetic relationship between isolates or their assignment to hybrid or parental classes; this is consistent with a highly dynamic chromosome complement in *Leishmania* promastigotes described both experimentally (Sterkers et al. 2011) and in field samples, for example in *L. donovani* in the Indian subcontinent (ISC; Imamura et al., 2016). Indeed, aneuploidy patterns do not seem to strongly segregate between Ethiopian *L. donovani* populations (Zackay et al. 2018) despite the degree of nucleotide diversity identified in these samples from one region of Ethiopia alone being much higher than in the ISC. For example, the average pair of samples in the main ISC population differ at only 88.3 sites whereas in the current cohort even two of the closely related ‘parent B’ isolates vary at an average of 1038 sites. Aneuploidy is known to be beneficial in allowing some single celled eukaryotes, for example *Saccharomyces* and *Candida*, to rapidly generate adaptive diversity (A. M. Selmecki et al. 2015), and likely contributes to adaptation in *Leishmania* (Mannaert et al. 2012; Prieto Barja et al. 2017). Aneuploidy is known to impact gene expression in *Leishmania* promastigotes (Dumetz et al. 2017; Iantorno et al. 2017). As *Leishmania* lacks classical regulation of transcription at initiation through promoters, this could contribute to parasite adaptation to at least some conditions (Laffitte et al. 2016; Mannaert et al. 2012). However, it is unclear how extensive aneuploidy variation in cultured promastigotes is relevant to the situation in either natural vectors, or in amastigotes *in vitro* or *in vivo:* while it is clear at least some variation does occur in both (Dumetz et al. 2017; Kumar et al. 2013) it appears to be much less widespread than in *in vitro* culture.

A striking result is that the sequence data suggests that isolate LdDM299, a recrudescent infection (from LdDM62) taken from the same patient was remarkably polysomic across all chromosomes relative to other isolates. Previous flow cytometry measurements of DNA content were suggestive of diploidy for both strains LdDM62 and LdDM299 and all other strains in the cohort (Gelanew et al. 2014) and are therefore incongruent. However a potential confounder was that the original cloned line that was sequenced was not available for cytometric analysis, with the isolate having undergone additional passages (>8). Somy in *Leishmania* can vary dramatically and rapidly in culture (Lachaud et al. 2014; Dumetz et al. 2017) and prolonged *in vitro* culture is known to systematically reduce ploidy in experimentally derived *T. cruzi* (Lewis et al. 2009). Here, relative somy between chromosomes is inferred from the coverage depth of reads mapped to each chromosome, while the baseline somy is determined from the allele frequency distribution. In principle, this could be misleading if the samples sequenced were mixtures of clones with many different somy levels, and single cell approaches such as FISH or single cell sequencing would be needed to fully disentangle this (Dujardin et al. 2014). However, the differences in allele frequency distributions between LdDM62 and LdDM299 for many chromosomes is particularly striking (Figure 2b), so there are at least genuine differences in the complement of chromosomes between the sequenced isolates. The consistently high dosage of some chromosomes – most strikingly chromosome 31 – are also broadly consistent with previous reports (Rogers et al. 2011; Downing et al. 2011; Dumetz et al. 2017). Together these provide reassurance that somy inferences are correct. We speculate that the apparent remarkable differences in somy between LdDM62 and LdDM299 isolated from the same HIV patient, could be an adaptive response to either chemotherapy or suppression of the patient’s immune response. SNP differences between LdM62 and LdM299 isolates were minimal, so aneuploidy variation could be a convenient mechanism to alter gene expression in response to drug pressure, as demonstrated in *Leishmania* (Mannaert et al. 2012) and conclusively in resistance of some pathogenic fungi to azole drugs (Kwon-Chung and Chang 2012; A. Selmecki, Forche, and Berman 2006).

Broadly, mitochondrial phylogenies corresponded to the expected nuclear genotypes (Figures 1 and 8 respectively). These data suggest hybrids uniparentally inherited parent type A mitochondrial genomes in agreement with inheritance patterns seen previously in other trypanosomatids (Messenger et al. 2012); (Satoskar and Snider 2009). There was some indication that LdDM62 and LdDM299, isolated from the same patient pre and post treatment, possessed some mitochondrial sequence diversity. However, bootstrap support was low. While all analyses support the clustering of LV9, parent A and hybrid isolates, the precise placement of different isolates within this cluster varied with details of the mitochondrial maxicircle assembly approach. In this context we do not interpret these small differences between nuclear and mitochondrial phylogenies as evidence of mitochondrial introgression. Mitochondrial introgression would be a very specific marker of hybridisation between populations and has been described in trypanosomatids, including *T. cruzi* (Messenger et al. 2012) and in many other organisms (Harrison and Larson 2014). Different ancestries between mitochondrial and nuclear genomes would not be expected between LdDM62 and the recrudescent infection LdDM299 in that they are likely to be the product of a single hybridisation event, based on near identical genomic structure and SNP profiles.

Current understanding regarding pattern and process of hybridisation in *Leishmania* is incomplete. Analysis of populations to detect and describe genomic variation in evolutionary recent hybrid isolates can confirm that hybridisation occurs in natural populations and provide insight into rates and patterns of recombination. For example, previous estimates based on genomic analysis form natural *L. infantum* isolates from Turkey indicate a hybridisation frequency of 1.3 x10^−5^ meioses per mitosis (Rogers et al. 2014). However characterisation of natural systems presents particular challenges: while co-localised isolates similar to the putative parents can sometimes be found, this is not guaranteed (Rogers et al. 2014). The number of independent meioses sampled in a natural population can be small and is consistently difficult to quantify. The recent ability to derive experimental hybrids in *L. major* (Akopyants et al. 2009) and now *L. donovani* (Sadlova et al. 2011); Yeo et al. unpublished data) can facilitate our understanding, as multiple replicated offspring from identical (and known) parents, frequency and distribution of cross-overs are easier to assess. Particular questions of interest might be to determine if recombination tends to occur at particular localised hot-spots? If so, are they associated with particular genomic features such as GC content or between polycistronic transcription units? Are crossing-over events associated with particularly high SNP mutation rates (Arbeithuber et al. 2015). In the current data we do not observe particular clusters of SNPs absent in the parental populations that could suggest this, as these data have limited power to detect these effects, which would require large number of observations of independent crossing-overs between the same parental haplotypes. Similarly, the contribution of gene conversion, often associated with meiosis, on either SNPs or tandem gene families are difficult to infer in these natural data. It is also important to note that ‘parental’ isolates represent here are only proxies for the true parents but results are strongly suggestive of multiple recombination events in Ethiopian *L. donovani* in recent evolutionary history. Encouragingly, experiments and subsequent derivation of experimental hybrids from phylogenetically similar parental genotypes also suggest frequent recombination in different sand fly vector species (Yeo et al, unpublished data). Experimental work will produce quantitative insights to support deeper understanding of the mechanisms and implications of recombination in *L. donovani* populations.

In conclusion we have presented genome-wide sequence data for putatively hybrid isolates of *L. donovani* from human VL cases in Ethiopia, together with isolates possessing putative parental like genotypes. We confirmed that 4 of the 5 putative hybrids are, indeed hybrid offspring derived from strains related to these parents, but the evolutionary history of these isolates is complex: representing at least 3 different histories. The haplotypic reconstructions, distribution of parent distinguishing SNPs and patterns of allele sharing are consistent with the occurence of more than one hybridisation event and/or intercrossing and backcrossing to parentals, which has not been observed in experimental crossing experiments to date. These data thus confirm the ability of *Leishmania* to hybridise extensively in natural populations. The population of *L. donovani* in Ethiopia has undergone multiple rounds of hybridisation, and we predict complex patterns of crossing would be revealed by a more substantial sample size. Together with progress in deriving experimental hybrids there is now promise of elucidating the mechanisms and other phylo-epidemiological aspects of recombination that have widespread implications regarding the spread, diagnosis and control of *L. donovani* populations.

## Materials and methods

We generated short-read paired-end sequence data for 11 isolates of *Leishmania donovani* from Ethiopia (see table 1). Full details of the origin and isolation of the strains used are described elsewhere (Gelanew et al. 2010, 2014). Briefly, all were visceral leishmania isolated between 2007 and 2009 from humans in northern Ethiopia. Of note, isolate DM299 was a relapse of DM62 isolated form a HIV infected patient post treatment (Libo Kemkem-Abdurafi).

DNA library preparation was performed by shearing genomic DNA into 400–600 base pair fragments by focused ultrasonication (Covaris Adaptive Focused Acoustics technology; AFA Inc., Woburn, USA), standard multiplex Illumina libraries were prepared using the NEBNext DNA Library Kit. The libraries were amplified with 8 cycles of PCR using Kapa HiFi DNA polymerase’ and were then pooled. 100bp paired-end reads were generated on the Illumina HiSeq 2000 v3 according to the manufacturer’s standard sequencing protocol. All sequencing data for these isolates are available from the ENA under project ERP106107. The LV9 strain (MHOM/ET/67/HU3 also known as MHOM/ET/67/L82) was originally isolated from a VL case in the Humera district in the far North of Ethiopia in 1967 (Bradley and Kirkley 1977). The JPCM5 strain (MCAN/ES/98/LLM877) is an *L. infantum* from Spain, isolated from a dog in 1998; BPK282 (MHOM/NP/03/BPK282/0cl2) was isolated from a human VL case in Nepal in 2003. Illumina whole-genome data for these isolates were obtained from the ENA database, with parasite material, sequencing approach and analysis of these data detailed in (Rogers et al. 2011) for JPCM5 and LV9, (Downing et al. 2011) for BPK282.

Reads for each isolate were mapped to the *L. donovani* LV9 reference assembly using SMALT v0.7.0.1 (Ponstigl 2010), indexing every second 13-mer (DePristo et al. 2011) and mapping repetitively with a minimum identity of 80% and maximum insert size of 1200bp, and mapping each read in the pair independently (*-x* flag). Variants were called using the HaplotypeCaller algorithm of Genome Analysis Toolkit v3.4 (DePristo et al. 2011), following best-practice guidelines (Van der Auwera et al. 2013) except as detailed below. Variant calls were first filtered to remove any overlapping with a mask generated with the GEM mappability tool (Marco-Sola et al. 2012) to identify non-unique 100bp sequences and to remove 100bp either side of any gaps within scaffolds. Subsequent filtering with the Genome Analysis Toolkit removed sites using the filtering parameters: DP >= 5*ploidy, DP <= 1.75*(chromosome median read depth), FS <= 13.0 or missing, SOR <= 3.0 or missing, ReadPosRankSum <= 3.1 AND ReadPosRankSum >= −3.1, BaseQRankSum <= 3.1 AND BaseQRankSum >= −3.1, MQRankSum <= 3.1 AND MQRankSum >= −3.1, ClippingRankSum <= 3.1 AND ClippingRankSum >= −3.1. Calls were made both assuming diploid genotypes for every chromosome across isolates, and using a somy estimated for each chromosome independently for each isolates. Somy was estimated using the EM approach described previously (Iantorno et al. 2017), and values checked by manual inspection of read depth and allele frequency data.

The whole-genome phylogeny and principal components analysis presented here were generated by using VCFtools v0.1.15 (Danecek et al. 2011) to convert the variants from GATK vcf format to the input format for plink, and then plink v1.90b3v (Purcell et al. 2007) was used for the principal components analysis and to generate pairwise distances (1 - identity by similarity). The pairwise distances were used to calculate a neighbour-joining phylogeny using the neighbor program from phylip v3.6.9 (Felsenstein 2005). Phasing was based on identifying illumina reads and read pairs linking heterozygous sites within each isolate, using the phase command in samtools v.0.1.19-44428cd (Li et al. 2009), with a block size (k) of 15; the phasing results did not differ for other values of k tested (11, 13, or 20) except k=30, where few variants were phased and no blocks > 1kb were shared by all isolates. Note that this phasing approach identifies heterozygous sites *de novo* from read mapping data rather than using the variant calls, and reconstructs at most two haplotypes at any locus. Phylogenies for the inferred haplotypes were generated using raxmlHPC v8.2.8 (Stamatakis 2014) under a GTR+I+G model of nucleotide substitution and otherwise default parameters.

Parent-distinguishing sites were identified as those for which both parent A isolates shared an identical homozygous genotype and all four parent B isolates were homozygous for a different allele. These sites could be unambiguously assigned as being derived from one or other parent in the putative hybrid isolates, assuming these other isolates were hybrids of these parents. To extend this analysis to other sites across the genome, a Hidden Markov model (HMM) was used to classify every 100bp window along the genome of the 5 suspected hybrid isolates by likely ancestry. Three hidden ancestry states (homozygous parent A, homozygous parent B and heterozygous from each parent) were used to explain the pattern across the genome of 4 observed parent-distinguishing SNP “symbols” (homozygous A, homozygous B, heterozygous and a non-determinate symbol for windows with either no parent-distinguishing SNPs or more than one state). The 100bp window size was chosen to make the HMM computationally tractable and so that almost every window (198,679 out of 202,940 across 5 isolates) was unambiguous for the observed symbol. All transitions between hidden states were allowed, but each hidden state could emit only the corresponding observation or the non-determinate symbol. Initial transition and emission probabilities and trained parameters are shown in Supplementary table 2; the trained parameters did not depend strongly on the initial parameters. The HMM was trained independently on each chromosome and isolate, and then average transition and emission parameters, weighted by chromosome lengths used to infer hidden states. Training and Viterbi decoding of the HMM was performed using the HMM package in R v3.3.0 (R core team 2016).

kDNA maxicircle genome sequences were generated by mapping illumina sequence data against the available maxicircle sequence assembly for *L. tarentolae* (Simpson et al. 1987) and using MITObim (Hahn, Bachmann, and Chevreux 2013) to perform iterative guided assembly with block size (k parameter) of 61 and with read trimming. This produced assemblies of between 19,611bp and 21,682bp in a single contig in each isolate (the *L. tarentolae* maxicircle is 20,992bp), including the entire transcribed region: tests using less strict criteria for assembly produced longer but less reliable assemblies. The assembled contigs were then rotated using CSA (Fernandes, Pereira, and Freitas 2009) before aligning with MAFFT v7.205 (Katoh and Standley 2013) with automated algorithm choice (--automated1 flag); the alignment was then trimmed with trimAl v1.4 (Fernandes, Pereira, and Freitas 2009; Capella-Gutiérrez, Silla-Martínez, and Gabaldón 2009). A maximum-likelihood phylogeny was inferred using raxmlHPC v8.2.8 under a GTR+I+G model of nucleotide substitutions (Stamatakis 2014) with 10 random addition-sequence replicates, and confidence in branches of the tree assessed with 500 bootstrap replicates.

## Acknowledgements

We thank the Wellcome Sanger Institute staff of the DNA pipelines at WSI for sequencing and generating sequencing libraries.

---

**Supplementary table 1.** Large (> 100bp) Structural variation between isolates (* this does not include LV9).

|  | Number of variants called | variants segregating in recent Ethiopian isolates* | Average heterozygosity in parent A isolates | Average heterozygosity in parent B isolates | Average heterozygosity in putative hybrids | Heterozygosity of Ld481 | Average heterozygosity of outgroups |
| --- | --- | --- | --- | --- | --- | --- | --- |
| Duplications | 169 | 95 | 0.64 | 0.60 | 0.68 | 0.69 | 0.15 |
| Deletions | 368 | 279 | 0.61 | 0.49 | 0.62 | 0.60 | 0.23 |
| Inversions | 282 | 147 | 0.63 | 0.56 | 0.69 | 0.65 | 0.17 |
| Insertions | 1 | 0 | 0 | 0 | 0 | 0 | 0 |
| Translocations | 264 | 123 | 0.44 | 0.36 | 0.42 | 0.44 | 0.16 |

**Supplementary table 2.** Initial transition (a) and emission (b) probability matrix and trained transition (c) and emission (d) probabilities for HMM. NA represents “Not Allowed” emissions from that state.

| From \ To | A | B | Het |
| --- | --- | --- | --- |
| A | 0.8 | 0.1 | 0.1 |
| B | 0.1 | 0.8 | 0.1 |
| Het | 0.05 | 0.05 | 0.9 |

(b)
| State \ Symbol | A | B | Het | Non-determinate |
| --- | --- | --- | --- | --- |
| A | 0.46 | NA | NA | 0.54 |
| B | NA | 0.46 | NA | 0.54 |
| Het | NA | NA | 0.46 | 0.54 |

(c)
| From \ To | A | B | Het |
| --- | --- | --- | --- |
| A | 0.94105911 | 0.03861683 | 0.02032406 |
| B | 0.08776222 | 0.88229226 | 0.02994551 |
| Het | 0.04503840 | 0.04030465 | 0.91465696 |

(d)
| State \ Symbol | A | B | Het | Non-determinate |
| --- | --- | --- | --- | --- |
| A | 0.1592378 | NA | NA | 0.8407622 |
| B | NA | 0.1653358 | NA | 0.8346642 |
| Het | NA | NA | 0.0878155 | 0.9121845 |

**Supplementary figure 1.**
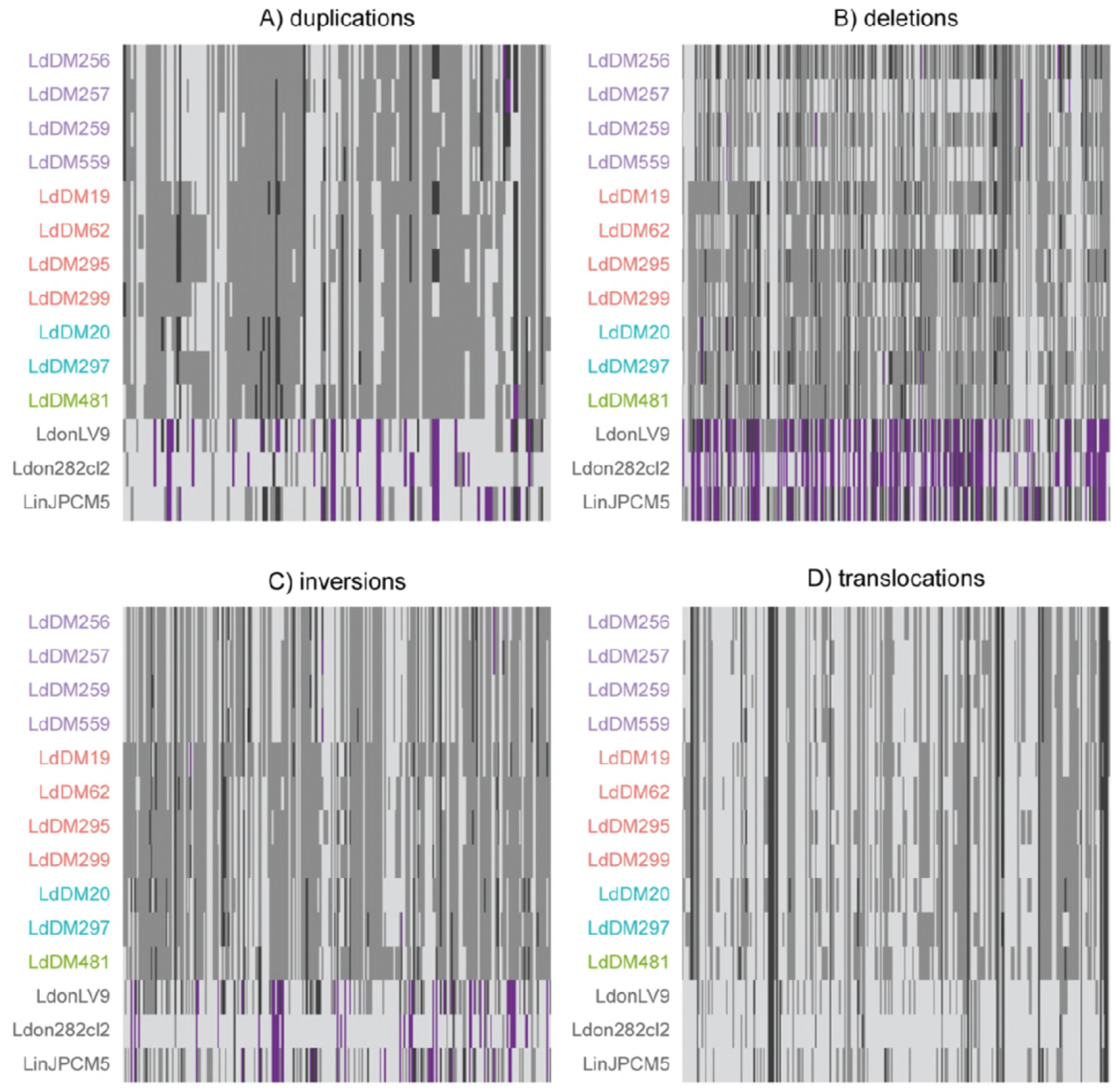
Overlaid DNA histograms for selected cloned *Leishmania* strains illustrating comparable (2*n*) DNA content representative of all hybrids (grey histogram) and both parental groups (cyan and red histogram). Gates were created for G1-0 (2*n*) peaks and for G2-M (4*n*) peaks. Each strain was tested in triplicate at a minimum and a control *Leishmania* strain was included in each run as an internal standard. Relative DNA content values were calculated as a ratio compared with the internal standard. Mean G1-0 values were taken to infer relative DNA content. The x-axes represent fluorescence intensity (arbitrary units) and the y-axes represent number of events in each channel.

## References

Akhoundi, Mohammad, Tim Downing, Jan Votýpka, Katrin Kuhls, Julius Lukeš, Arnaud Cannet, Christophe Ravel, et al. 2017. “Leishmania Infections: Molecular Targets and Diagnosis.” Molecular Aspects of Medicine 57 (October): 1–29.

Akhoundi, Mohammad, Katrin Kuhls, Arnaud Cannet, Jan Votýpka, Pierre Marty, Pascal Delaunay, and Denis Sereno. 2016. “A Historical Overview of the Classification, Evolution, and Dispersion of Leishmania Parasites and Sandflies.” PLoS Neglected Tropical Diseases 10 (3): e0004349.

Akopyants, Natalia S., Nicola Kimblin, Nagila Secundino, Rachel Patrick, Nathan Peters, Phillip Lawyer, Deborah E. Dobson, Stephen M. Beverley, and David L. Sacks. 2009. “Demonstration of Genetic Exchange during Cyclical Development of Leishmania in the Sand Fly Vector.” Science 324 (5924): 265–68.

Alvar, Jorge, Iván D. Vélez, Caryn Bern, Mercé Herrero, Philippe Desjeux, Jorge Cano, Jean Jannin, Margriet den Boer, and WHO Leishmaniasis Control Team. 2012. “Leishmaniasis Worldwide and Global Estimates of Its Incidence.” PloS One 7 (5): e35671.

Arbeithuber, Barbara, Andrea J. Betancourt, Thomas Ebner, and Irene Tiemann-Boege. 2015. “Crossovers Are Associated with Mutation and Biased Gene Conversion at Recombination Hotspots.” Proceedings of the National Academy of Sciences of the United States of America 112 (7): 2109–14.

Bradley, D. J., and J. Kirkley. 1977. “Regulation of Leishmania Populations within the Host. I. the Variable Course of Leishmania Donovani Infections in Mice.” Clinical and Experimental Immunology 30 (1): 119–29.

Capella-Gutiérrez, Salvador, José M. Silla-Martínez, and Toni Gabaldón. 2009. “trimAl: A Tool for Automated Alignment Trimming in Large-Scale Phylogenetic Analyses.” Bioinformatics 25 (15): 1972–73.

Danecek, Petr, Adam Auton, Goncalo Abecasis, Cornelis A. Albers, Eric Banks, Mark A. DePristo, Robert E. Handsaker, et al. 2011. “The Variant Call Format and VCFtools.” Bioinformatics 27 (15): 2156–58.

DePristo, Mark A., Eric Banks, Ryan Poplin, Kiran V. Garimella, Jared R. Maguire, Christopher Hartl, Anthony A. Philippakis, et al. 2011. “A Framework for Variation Discovery and Genotyping Using next-Generation DNA Sequencing Data.” Nature Genetics 43 (5): 491–98.

Desjardins, Christopher A., Charles Giamberardino, Sean M. Sykes, Chen-Hsin Yu, Jennifer L. Tenor, Yuan Chen, Timothy Yang, et al. 2017. “Population Genomics and the Evolution of Virulence in the Fungal Pathogen Cryptococcus Neoformans.” Genome Research 27 (7): 1207–19.

Downing, Tim, Hideo Imamura, Saskia Decuypere, Taane G. Clark, Graham H. Coombs, James A. Cotton, James D. Hilley, et al. 2011. “Whole Genome Sequencing of Multiple Leishmania Donovani Clinical Isolates Provides Insights into Population Structure and Mechanisms of Drug Resistance.” Genome Research 21 (12): 2143–56.

Dujardin, Jean-Claude, An Mannaert, Caroline Durrant, and James A. Cotton. 2014. “Mosaic Aneuploidy in Leishmania: The Perspective of Whole Genome Sequencing.” Trends in Parasitology 30 (12): 554–55.

Dumetz, F., H. Imamura, M. Sanders, V. Seblova, J. Myskova, P. Pescher, M. Vanaerschot, et al. 2017. “Modulation of Aneuploidy in Leishmania Donovani during Adaptation to Different In Vitro and In Vivo Environments and Its Impact on Gene Expression.” mBio 8 (3). https://doi.org/10.1128/mBio.00599-17.

Ene, Iuliana V., and Richard J. Bennett. 2014. “The Cryptic Sexual Strategies of Human Fungal Pathogens.” Nature Reviews. Microbiology 12 (4): 239–51.

Felsenstein, J. 2005. “PHYLIP (Phylogeny Inference Package) Version 3.6.”

Fernandes, Francisco, Luísa Pereira, and Ana T. Freitas. 2009. “CSA: An Efficient Algorithm to Improve Circular DNA Multiple Alignment.” BMC Bioinformatics 10 (July): 230.

Gelanew, Tesfaye, Asrat Hailu, Gabriele Schőnian, Michael D. Lewis, Michael A. Miles, and Matthew Yeo. 2014. “Multilocus Sequence and Microsatellite Identification of Intra-Specific Hybrids and Ancestor-like Donors among Natural Ethiopian Isolates of Leishmania Donovani.” International Journal for Parasitology 44 (10): 751–57.

Gelanew, Tesfaye, Katrin Kuhls, Zewdu Hurissa, Teklu Weldegebreal, Workagegnehu Hailu, Aysheshm Kassahun, Tamrat Abebe, Asrat Hailu, and Gabriele Schönian. 2010. “Inference of Population Structure of Leishmania Donovani Strains Isolated from Different Ethiopian Visceral Leishmaniasis Endemic Areas.” PLoS Neglected Tropical Diseases 4 (11): e889.

Hahn, Christoph, Lutz Bachmann, and Bastien Chevreux. 2013. “Reconstructing Mitochondrial Genomes Directly from Genomic next-Generation Sequencing Reads--a Baiting and Iterative Mapping Approach.” Nucleic Acids Research 41 (13): e129.

Hamad, S. H., Ahmed M. Musa, Eltahir A. G. Khalil, Tamrat Abebe, Brima M. Younis, Mona E. E. Elthair, Ahmed M. EL-Hassan, Asrat Hailu, and Aldert Bart. 2011. “Leishmania: Probable Genetic Hybrids between Species in Sudanese Isolates.” Journal of Microbiology and Antimicrobials 3: 142–45.

Harrison, Richard G., and Erica L. Larson. 2014. “Hybridization, Introgression, and the Nature of Species Boundaries.” The Journal of Heredity 105 Suppl 1: 795–809.

Herricks, Jennifer R., Peter J. Hotez, Valentine Wanga, Luc E. Coffeng, Juanita A. Haagsma, María-Gloria Basáñez, Geoffrey Buckle, et al. 2017. “The Global Burden of Disease Study 2013: What Does It Mean for the NTDs?” PLoS Neglected Tropical Diseases 11 (8): e0005424.

Herwaldt, B. L. 1999. “Leishmaniasis.” The Lancet 354 (9185): 1191–99.

Iantorno, Stefano A., Caroline Durrant, Asis Khan, Mandy J. Sanders, Stephen M. Beverley, Wesley C. Warren, Matthew Berriman, David L. Sacks, James A. Cotton, and Michael E. Grigg. 2017. “Gene Expression in Leishmania Is Regulated Predominantly by Gene Dosage.” mBio 8 (5). https://doi.org/10.1128/mBio.01393-17.

Inbar, Ehud, Natalia S. Akopyants, Melanie Charmoy, Audrey Romano, Phillip Lawyer, Dia-Eldin A. Elnaiem, Florence Kauffmann, et al. 2013. “The Mating Competence of Geographically Diverse Leishmania Major Strains in Their Natural and Unnatural Sand Fly Vectors.” PLoS Genetics 9 (7): e1003672.

Jensen, Robert E., and Paul T. Englund. 2012. “Network News: The Replication of Kinetoplast DNA.” Annual Review of Microbiology 66: 473–91.

Katoh, Kazutaka, and Daron M. Standley. 2013. “MAFFT Multiple Sequence Alignment Software Version 7: Improvements in Performance and Usability.” Molecular Biology and Evolution 30 (4): 772–80.

Kuhls, Katrin, Elisa Cupolillo, Soraia O. Silva, Carola Schweynoch, Mariana Côrtes Boité, Maria N. Mello, Isabel Mauricio, Michael Miles, Thierry Wirth, and Gabriele Schönian. 2013. “Population Structure and Evidence for Both Clonality and Recombination among Brazilian Strains of the Subgenus Leishmania (Viannia).” PLoS Neglected Tropical Diseases 7 (10): e2490.

Kuhls, Katrin, Lyvia Keilonat, Sebastian Ochsenreither, Matthias Schaar, Carola Schweynoch, Wolfgang Presber, and Gabriele Schönian. 2007. “Multilocus Microsatellite Typing (MLMT) Reveals Genetically Isolated Populations between and within the Main Endemic Regions of Visceral Leishmaniasis.” Microbes and Infection / Institut Pasteur 9 (3): 334–43.

Kumar, Pranav, Robert Lodge, Frédéric Raymond, Jean-François Ritt, Pascal Jalaguier, Jacques Corbeil, Marc Ouellette, and Michel J. Tremblay. 2013. “Gene Expression Modulation and the Molecular Mechanisms Involved in Nelfinavir Resistance in Leishmania Donovani Axenic Amastigotes.” Molecular Microbiology 89 (3): 565–82.

Kwon-Chung, Kyung J., and Yun C. Chang. 2012. “Aneuploidy and Drug Resistance in Pathogenic Fungi.” PLoS Pathogens 8 (11): e1003022.

Lachaud, Laurence, Nathalie Bourgeois, Nada Kuk, Christelle Morelle, Lucien Crobu, Gilles Merlin, Patrick Bastien, Michel Pagès, and Yvon Sterkers. 2014. “Constitutive Mosaic Aneuploidy Is a Unique Genetic Feature Widespread in the Leishmania Genus.” Microbes and Infection / Institut Pasteur 16 (1): 61–66.

Laffitte, Marie-Claude N., Philippe Leprohon, Barbara Papadopoulou, and Marc Ouellette. 2016. “Plasticity of the Leishmania Genome Leading to Gene Copy Number Variations and Drug Resistance.” F1000Research 5 (September): 2350.

Lewis, Michael D., Martin S. Llewellyn, Michael W. Gaunt, Matthew Yeo, Hernán J. Carrasco, and Michael A. Miles. 2009. “Flow Cytometric Analysis and Microsatellite Genotyping Reveal Extensive DNA Content Variation in Trypanosoma Cruzi Populations and Expose Contrasts between Natural and Experimental Hybrids.” International Journal for Parasitology 39 (12): 1305–17.

Li, Heng, Bob Handsaker, Alec Wysoker, Tim Fennell, Jue Ruan, Nils Homer, Gabor Marth, Goncalo Abecasis, Richard Durbin, and 1000 Genome Project Data Processing Subgroup. 2009. “The Sequence Alignment/Map Format and SAMtools.” Bioinformatics 25 (16): 2078–79.

Mannaert, An, Tim Downing, Hideo Imamura, and Jean-Claude Dujardin. 2012. “Adaptive Mechanisms in Pathogens: Universal Aneuploidy in Leishmania.” Trends in Parasitology 28 (9): 370–76.

Marco-Sola, Santiago, Michael Sammeth, Roderic Guigó, and Paolo Ribeca. 2012. “The GEM Mapper: Fast, Accurate and Versatile Alignment by Filtration.” Nature Methods 9 (12): 1185–88.

McMullan, Mark, Anastasia Gardiner, Kate Bailey, Eric Kemen, Ben J. Ward, Volkan Cevik, Alexandre Robert-Seilaniantz, et al. 2015. “Evidence for Suppression of Immunity as a Driver for Genomic Introgressions and Host Range Expansion in Races of Albugo Candida, a Generalist Parasite.” eLife 4 (February). https://doi.org/10.7554/eLife.04550.

Messenger, Louisa A., Martin S. Llewellyn, Tapan Bhattacharyya, Oscar Franzén, Michael D. Lewis, Juan David Ramírez, Hernan J. Carrasco, Björn Andersson, and Michael A. Miles. 2012. “Multiple Mitochondrial Introgression Events and Heteroplasmy in Trypanosoma Cruzi Revealed by Maxicircle MLST and Next Generation Sequencing.” PLoS Neglected Tropical Diseases 6 (4): e1584.

Nolder, Debbie, Norma Roncal, Clive R. Davies, Alejandro Llanos-Cuentas, and Michael A. Miles. 2007. “Multiple Hybrid Genotypes of Leishmania (viannia) in a Focus of Mucocutaneous Leishmaniasis.” The American Journal of Tropical Medicine and Hygiene 76 (3): 573–78.

Peacock, Lori, Mick Bailey, Mark Carrington, and Wendy Gibson. 2014. “Meiosis and Haploid Gametes in the Pathogen Trypanosoma Brucei.” Current Biology: CB 24 (2): 181–86.

Peacock, Lori, Mick Bailey, and Wendy Gibson. 2016. “Dynamics of Gamete Production and Mating in the Parasitic Protist Trypanosoma Brucei.” Parasites & Vectors 9 (1). https://doi.org/10.1186/s13071-016-1689-9.

Ponstigl, H. 2010. “Smalt.” 2010. https://sourceforge.net/projects/smalt/.

Prieto Barja, Pablo, Pascale Pescher, Giovanni Bussotti, Franck Dumetz, Hideo Imamura, Darek Kedra, et al. 2017. “Haplotype Selection as an Adaptive Mechanism in the Protozoan Pathogen Leishmania Donovani.” Nature Ecology & Evolution 1 (12): 1961–69.

Pugach, Irina, Rostislav Matveyev, Andreas Wollstein, Manfred Kayser, and Mark Stoneking. 2011. “Dating the Age of Admixture via Wavelet Transform Analysis of Genome-Wide Data.” Genome Biology 12 (2): R19.

Purcell, Shaun, Benjamin Neale, Kathe Todd-Brown, Lori Thomas, Manuel A. R. Ferreira, David Bender, Julian Maller, et al. 2007. “PLINK: A Tool Set for Whole-Genome Association and Population-Based Linkage Analyses.” American Journal of Human Genetics 81 (3): 559–75.

Ramírez, Juan David, and Martin S. Llewellyn. 2014. “Reproductive Clonality in Protozoan Pathogens-Truth or Artefact?” Molecular Ecology 23 (17): 4195–4202.

Ravel, Christophe, Sofia Cortes, Francine Pratlong, Florent Morio, Jean-Pierre Dedet, and Lenea Campino. 2006. “First Report of Genetic Hybrids between Two Very Divergent Leishmania Species: Leishmania Infantum and Leishmania Major.” International Journal for Parasitology 36 (13): 1383–88.

R core team. 2016. R: A Language and Environment for Statistical Computing. https://www.R-project.org/.

Rogers, Matthew B., Tim Downing, Barbara A. Smith, Hideo Imamura, Mandy Sanders, Milena Svobodova, Petr Volf, Matthew Berriman, James A. Cotton, and Deborah F. Smith. 2014. “Genomic Confirmation of Hybridisation and Recent Inbreeding in a Vector-Isolated Leishmania Population.” PLoS Genetics 10 (1): e1004092.

Rogers, Matthew B., James D. Hilley, Nicholas J. Dickens, Jon Wilkes, Paul A. Bates, Daniel P. Depledge, David Harris, et al. 2011. “Chromosome and Gene Copy Number Variation Allow Major Structural Change between Species and Strains of Leishmania.” Genome Research 21 (12): 2129–42.

Romano, Audrey, Ehud Inbar, Alain Debrabant, Melanie Charmoy, Phillip Lawyer, Flavia Ribeiro-Gomes, Mourad Barhoumi, et al. 2014. “Cross-Species Genetic Exchange between Visceral and Cutaneous Strains of Leishmania in the Sand Fly Vector.” Proceedings of the National Academy of Sciences of the United States of America 111 (47): 16808–13.

Ropars, Jeanne, Corinne Maufrais, Dorothée Diogo, Marina Marcet-Houben, Aurélie Perin, Natacha Sertour, Kevin Mosca, et al. 2018. “Gene Flow Contributes to Diversification of the Major Fungal Pathogen Candida Albicans.” Nature Communications 9 (1): 2253.

Rougeron, Virginie, Thierry De Meeûs, Mallorie Hide, Etienne Waleckx, Herman Bermudez, Jorge Arevalo, Alejandro Llanos-Cuentas, et al. 2009. “Extreme Inbreeding in Leishmania Braziliensis.” Proceedings of the National Academy of Sciences of the United States of America 106 (25): 10224–29.

Rougeron, Virginie, Thierry De Meeûs, Sandrine Kako Ouraga, Mallorie Hide, and Anne-Laure Bañuls. 2010. “‘Everything You Always Wanted to Know about Sex (but Were Afraid to Ask)’ in Leishmania after Two Decades of Laboratory and Field Analyses.” PLoS Pathogens 6 (8): e1001004.

Sadlova, Jovana, Matthew Yeo, Veronika Seblova, Michael D. Lewis, Isabel Mauricio, Petr Volf, and Michael A. Miles. 2011. “Visualisation of Leishmania Donovani Fluorescent Hybrids during Early Stage Development in the Sand Fly Vector.” PloS One 6 (5): e19851.

Satoskar, Abhay, and Heidi Snider. 2009. “Faculty of 1000 Evaluation for Demonstration of Genetic Exchange during Cyclical Development of Leishmania in the Sand Fly Vector.” F1000 - Post-Publication Peer Review of the Biomedical Literature. https://doi.org/10.3410/f.1159113.621459.

Schönian, Gabriele, Elisa Cupolillo, and Isabel Mauricio. 2012. “Molecular Evolution and Phylogeny of Leishmania.” In Drug Resistance in Leishmania Parasites, 15–44.

Schönian, G., K. Kuhls, and I. L. Mauricio. 2011. “Molecular Approaches for a Better Understanding of the Epidemiology and Population Genetics of Leishmania.” Parasitology 138 (4): 405–25.

Schwenkenbecher, Jan M., Thierry Wirth, Lionel F. Schnur, Charles L. Jaffe, Henk Schallig, Amer Al-Jawabreh, Omar Hamarsheh, Kifaya Azmi, Francine Pratlong, and Gabriele Schönian. 2006. “Microsatellite Analysis Reveals Genetic Structure of Leishmania Tropica.” International Journal for Parasitology 36 (2): 237–46.

Seblova, Veronika, Jovana Sadlova, Barbora Vojtkova, Jan Votypka, Simon Carpenter, Paul Andrew Bates, and Petr Volf. 2015. “The Biting Midge Culicoides Sonorensis (Diptera: Ceratopogonidae) Is Capable of Developing Late Stage Infections of Leishmania Enriettii.” PLoS Neglected Tropical Diseases 9 (9): e0004060.

Seblova, Veronika, Vera Volfova, Vit Dvorak, Katerina Pruzinova, Jan Votypka, Aysheshm Kassahun, Teshome Gebre-Michael, Asrat Hailu, Alon Warburg, and Petr Volf. 2013. “Phlebotomus Orientalis Sand Flies from Two Geographically Distant Ethiopian Localities: Biology, Genetic Analyses and Susceptibility to Leishmania Donovani.” PLoS Neglected Tropical Diseases 7 (4): e2187.

Selmecki, Anna, Anja Forche, and Judith Berman. 2006. “Aneuploidy and Isochromosome Formation in Drug-Resistant Candida Albicans.” Science 313 (5785): 367–70.

Selmecki, Anna M., Yosef E. Maruvka, Phillip A. Richmond, Marie Guillet, Noam Shoresh, Amber L. Sorenson, Subhajyoti De, et al. 2015. “Polyploidy Can Drive Rapid Adaptation in Yeast.” Nature 519 (7543): 349–52.

Simpson, L., N. Neckelmann, V. F. de la Cruz, A. M. Simpson, J. E. Feagin, D. P. Jasmer, and K. Stuart. 1987. “Comparison of the Maxicircle (mitochondrial) Genomes of Leishmania Tarentolae and Trypanosoma Brucei at the Level of Nucleotide Sequence.” The Journal of Biological Chemistry 262 (13): 6182–96.

Stamatakis, Alexandros. 2014. “RAxML Version 8: A Tool for Phylogenetic Analysis and Post-Analysis of Large Phylogenies.” Bioinformatics 30 (9): 1312–13.

Sterkers, Yvon, Lucien Crobu, Laurence Lachaud, Michel Pagès, and Patrick Bastien. 2014. “Parasexuality and Mosaic Aneuploidy in Leishmania: Alternative Genetics.” Trends in Parasitology 30 (9): 429–35.

Sterkers, Yvon, Laurence Lachaud, Lucien Crobu, Patrick Bastien, and Michel Pagès. 2011. “FISH Analysis Reveals Aneuploidy and Continual Generation of Chromosomal Mosaicism in Leishmania Major.” Cellular Microbiology 13 (2): 274–83.

Tibayrenc, M., F. Kjellberg, and F. J. Ayala. 1990. “A Clonal Theory of Parasitic Protozoa: The Population Structures of Entamoeba, Giardia, Leishmania, Naegleria, Plasmodium, Trichomonas, and Trypanosoma and Their Medical and Taxonomical Consequences.” Proceedings of the National Academy of Sciences of the United States of America 87 (7): 2414–18.

Turner, C. M. R., G. Hide, N. Buchanan, and A. Tait. 1995. “Trypanosoma Brucei: Inheritance of Kinetoplast DNA Maxicircles in a Genetic Cross and Their Segregation during Vegetative Growth.” Experimental Parasitology 80 (2): 234–41.

Twyford, A. D., and R. A. Ennos. 2011. “Next-Generation Hybridization and Introgression.” Heredity 108 (3): 179–89.

Van der Auwera, Geraldine A., Mauricio O. Carneiro, Christopher Hartl, Ryan Poplin, Guillermo del Angel, Ami Levy-Moonshine, Tadeusz Jordan, et al. 2013. “From FastQ Data to High-Confidence Variant Calls: The Genome Analysis Toolkit Best Practices Pipeline.” In Current Protocols in Bioinformatics, 11.10.1–11.10.33.

Volf, Petr, Ivana Benkova, Jitka Myskova, Jovana Sadlova, Lenea Campino, and Christophe Ravel. 2007. “Increased Transmission Potential of Leishmania major/Leishmania Infantum Hybrids.” International Journal for Parasitology 37 (6): 589–93.

Zackay, Arie, James A. Cotton, Mandy Sanders, Asrat Hailu, Abedelmajeed Nasereddin, Alon Warburg, and Charles L. Jaffe. 2018. “Genome Wide Comparison of Ethiopian Leishmania Donovani Strains Reveals Differences Potentially Related to Parasite Survival.” PLoS Genetics 14 (1): e1007133.

